# Neural-specific alterations in glycosphingolipid biosynthesis and cell signaling associated with two human ganglioside GM3 Synthase Deficiency variants

**DOI:** 10.1101/2021.07.29.454399

**Authors:** Michelle Dookwah, Shannon K. Wagner, Mayumi Ishihara, Seok-Ho Yu, Heidi Ulrichs, Michael J. Kulik, Nadja Zeltner, Stephen Dalton, Kevin A. Strauss, Kazuhiro Aoki, Richard Steet, Michael Tiemeyer

## Abstract

GM3 Synthase Deficiency (GM3SD) is a neurodevelopmental disorder resulting from pathogenic variants in the *ST3GAL5* gene, which encodes GM3 synthase, a glycosphingolipid (GSL)-specific sialyltransferase. This enzyme adds a single *α*3-linked sialic acid to the terminal galactose of lactosylceramide (LacCer) to produce the monosialylated ganglioside GM3. In turn, GM3 is extended by other glycosyltransferases to generate nearly all the complex gangliosides enriched in neural tissue. Pathogenic mechanisms that account for neural phenotypes associated with GM3SD are not known. To explore how loss of GM3 impacts neural-specific glycolipid glycosylation and cell signaling, GM3SD patient fibroblasts bearing one of two different *ST3GAL5* variants were reprogrammed to induced pluripotent stem cells (iPSCs) and then differentiated to neural crest cells (NCCs). GM3 and GM3-derived gangliosides were undetectable in iPSCs and NCCs from both variants, while LacCer precursor levels were elevated compared to wildtype (WT). NCCs of both variants synthesized elevated levels of neutral lacto- and globo-series, as well as minor alternatively sialylated, GSLs compared to WT. Shifts in ceramide profiles associated with iPSC and NCC GSLs were also detected in GM3SD variants. Altered GSL profiles in the GM3SD cells were accompanied by dynamic changes in the cell surface proteome, protein O-GlcNAcylation, and receptor tyrosine kinase abundance. GM3SD cells also exhibited increased apoptosis and sensitivity to erlotnib, an inhibitor of epidermal growth factor receptor signaling. Pharmacologic inhibition of O-GlcNAcase increased protein O-GlcNAcylation and significantly rescued baseline and erlotnib-induced apoptosis. Collectively, these findings indicate broad effects on cell signaling during differentiation of GM3SD patient-derived iPSCs to NCCs. Thus, human GM3SD cells provide a novel platform to investigate structure/function relationships that connect GSL diversity to cell signaling, cell survival, and neural differentiation.

## INTRODUCTION

GM3 Synthase Deficiency (GM3SD) is an autosomal recessive disorder characterized by infantile-onset epileptic encephalopathy, global developmental delay, postnatal microcephaly, congenital hearing loss, cortical visual impairment, and hyper- and hypo-pigmented epidermis [1, 2]. GM3SD is caused by pathogenic variants in the *ST3GAL5* gene that encodes for the sialyltransferase known as GM3 synthase. ST3GAL5 catalyzes the production of the ganglioside GM3, which is the precursor for all complex a- b-, and c-type ganglioside glycosphingolipids (GSLs) in the human brain (Figure 1). Gangliosides account for >80% of brain glycans and have been implicated in neural-glial cell interactions, neuronal proliferation and survival, membrane microdomain formation, and cell signaling [3, 4]. The significant clinical consequences of GM3SD are not entirely unexpected, in light of the impact of targeted biosynthetic knockouts in mice, and of the accumulated data highlighting in vitro functions of gangliosides [5–7]. Nonetheless, the functional impact of ganglioside-deficiency on neural cell differentiation, signaling, and survival in the context of neural cells in culture has not been studied, leaving gaps in our understanding of the neuronal pathogenesis associated with GM3SD.

**Figure 1.**
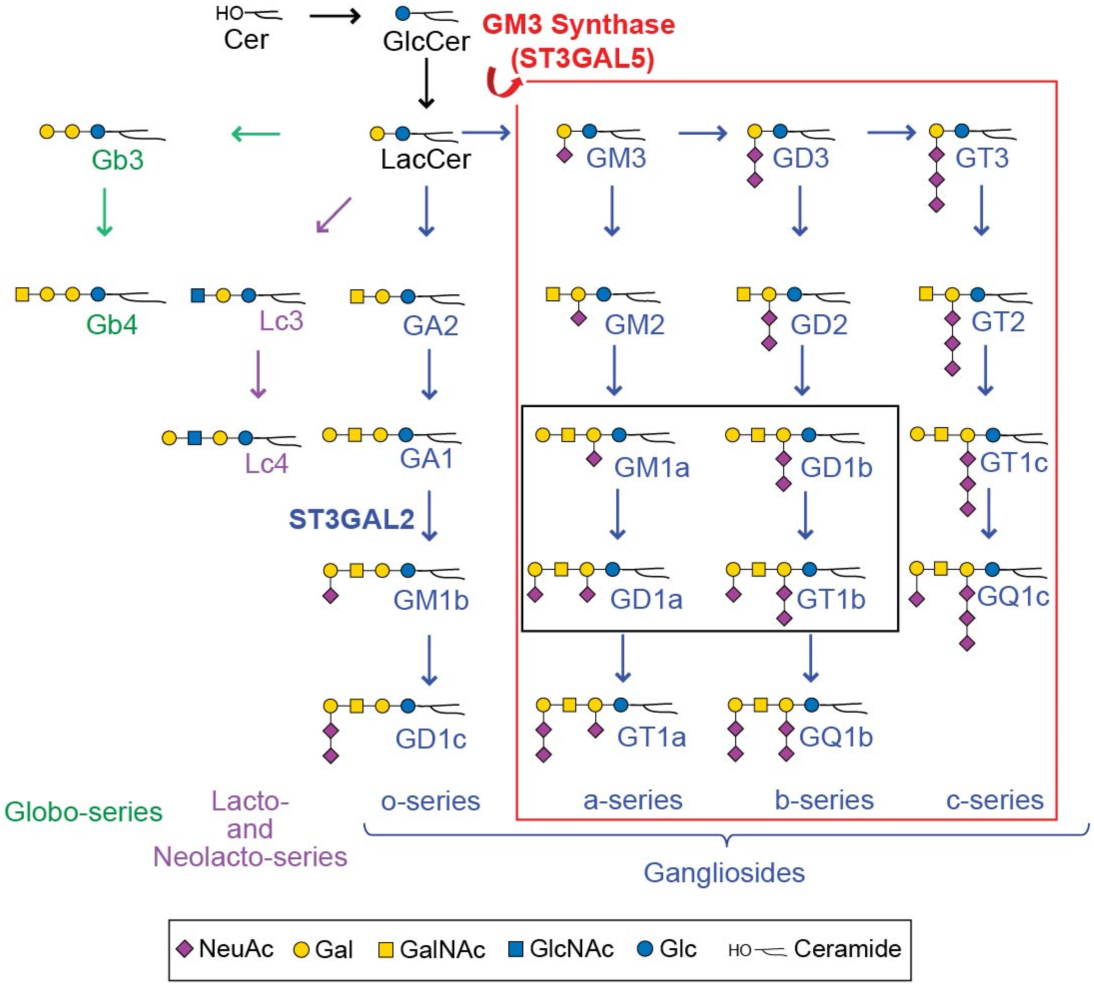
Pathways of human glycosphingolipid biosynthesis. Ceramide (Cer) is glucosylated to form glucosylceramide (GlcCer), which is elongated with a galactose to form lactosylceramide (LacCer). LacCer is the precursor for glycosphingolipid (GSL) elongation into multiple biosynthetic pathways, including the globo-, lacto/neolacto-, and ganglio-series. Sialylation of the Gal residue of LacCer by ST3GAL5 (GM3 Synthase) generates the simplest ganglio-series GSL, known as GM3. GM3 is the essential precursor for the production of all a-, b-, and c- series gangliosides (***red box***). Of these gangliosides, four are the most abundant GSLs found in neural tissue, GM1a, GD1a, GD1b and GT1b (***black box***). Graphic representations of GSL monosaccharide residues are consistent with Standard Nomenclature for Glycomics guidelines (SNFG).

GM3SD has been associated with several cohorts worldwide, exhibiting allelic diversity based in part on genetic ancestry. Salt & Pepper Syndrome was described as one of the first cohorts of GM3SD, affecting a single family in South Carolina, USA [2, 8]. The variant associated with Salt & Pepper Syndrome, *ST3GAL5* c.1063G>A (p.Glu355Lys), encodes for a mutant form of the enzyme lacking any enzymatic activity [9]. A more prevalent variant identified in the Amish population, c.862C>T (p.Arg288Ter), encodes for a truncated and inactive enzyme [9, 10]. The Amish variant is estimated to have a carrier frequency of 5-6% within this population, which translates to approximately 1 in 1200 births [1]. The p.Arg288Ter variant has also been reported in French and Pakistani cohorts [11, 12]. Additional variants, compound heterozygous ST3GAL5 mutations (p.Gly201Arg and p.Cys195Ser), were identified in two Korean siblings [13]. All of the currently identified and characterized pathogenic ST3GAL5 variants are null for enzymatic activity and share significant overlap in their clinical presentations [14].

To investigate the molecular and neural-specific changes resulting from loss of GM3, we generated induced pluripotent stem cells (iPSCs) from Salt & Pepper (p.Glu355Lys) and Amish cohort (p.Arg288Ter) fibroblasts. These cells were subsequently differentiated towards the neural crest cell (NCC) lineage, and the impact of GM3 synthase deficiency on neural-specific glycosphingolipid biosynthesis and cellular signaling responses was monitored across the differentiation time course. Neural crest was chosen as an initial lineage of study based on skin pigmentation phenotypes described in patients, high homogeneity of the NCC population derived from iPSCs, and the broad impact of neural crest derivatives on somatic function. This investigation identified several common GSL profile alterations in the two variants, including changes in ceramide structures and loss of complex gangliosides. These GSL changes were accompanied by broad fluctuations in multiple cell signaling pathways and an abnormal abundance of several lipid raft-associated cell surface proteins in the two patient lines. The implications of these findings for the neuronal pathogenesis of GM3SD and molecular phenotypes that distinguish the two patient lines are discussed. These results establish a baseline for the use of patient-derived cells for further investigating the impact of GSL diversity on cell signaling and differentiation in multiple tissue and cell-specific lineages.

## MATERIALS AND METHODS

### Generation of induced pluripotent stem cells

Wild-type iPSC lines hiPSK3 and hiC3 were, respectively, acquired from Steve Duncan, University of Wisconsin, or generated from ATCC CRL-1509 human fibroblasts [15]. Generation of p.Glu355Lys iPSCs (from ST3Gal5 c.1063G>A (p.Glu355Lys) patient fibroblasts) and p.Arg288Ter iPSCs (from ST3GAL5 c.862C>T (p.Arg288Ter) patient fibroblasts) was performed using the CytoTune-iPS Sendai Reprogramming Kit (Invitrogen, # A13780) and the CytoTune-iPS 2.0 Sendai Reprogramming Kit (ThermoFisher, # A16517), respectively, using the manufacturer’s conditions and as described previoulsy [16]. The p.Glu355Lys donor was a 13 year old male of African-American descent and the p.Arg288Ter donor was a 12 year old male and a member of the Amish community. A single iPSC clone was generated for the p.Glu355Lys variant and 3 clones were generated for the p.Arg288Ter variant. Each subclone was passaged at least 10 times to ensure removal of residual reprogramming virus before further characterization. Loss of virus was monitored by staining with anti-Sendai virus antibody (Invitrogen, cat #14649482).

iPSC clones were initially tested for pluripotency by assessing expression of pluripotency markers Oct4 (Cell Signaling, # C30A3), SSEA3/4 (Santa Cruz, # sc-21704), and Sox2 (R&D, # MAB2018), and by inducing differentiation into neuroectodermal cells [17]. For neuroectoderm differentiation, iPSCs were seeded at 260,000 cells/cm^2^ in 6 wells in Essential 8 media on matrigel. The next day, the media was changed to Essential 6 containing 100 nM LDN, 10 µM SB and 2 µM XAV until day 2 when the media was changed to Essential 6 containing 100 nM LDN, 10 µM SB. Cells were fixed and stained for PAX6 (Biolegend, # 901301/B201255) and DAPI at day 4 and exhibited efficient differentiation into the neuroectodermal lineage (**Supplement Figure 1**). All iPSCs were tested for mycoplasma every 2 weeks.

### Neural Crest differentiation of iPSCs

Patient and WT iPSCs were differentiated to neural crest cells (NCCs) according to Menendez, Kulik [16]. Briefly, a 60 mm dish of ∼90% confluent iPSCs was dissociated with Accutase (Innovative Cell Technologies, # AT104) and resuspended to a density of ∼9.2×10^4^ cells per cm^2^ on Geltrex-coated plates in neural crest media composed of DMEM/F12 without glutamine (Cellgro), 2% Probumin, Life Science Grade (Millipore), 1× Non-essential amino acids (Cellgro), 2 mM GlutaGro (Cellgro), 1× Antibiotic/Antimycotic (Cellgro), 10 µg/ml Human Transferrin, (Athens Research & Tech),1× Trace elements A, B, & C (Cellgro), 50 µg/ml L-Ascorbic acid 2-phosphate sesqimagnesium salt hydrate (Sigma), 10 ng/ml Heregulin β (Peprotech), 200 ng/ml Long R3-IGF1 (SAFC/Sigma), 8 ng/ml bFGF (RnD Systems), 2 µM GSK3 inhibitor IX (BIO) (Tocris), and 20 µM SB431542 (Tocris). Media was changed daily, and at confluence, cells were split 1:4-1:6 (vol/vol) onto Geltrex-coated plates. Cells exhibited NCC morphology between day 7-10 of differentiation. By Day 15, cells tested positive by immunohistochemistry for NC markers HNK1 (Sigma Aldrich, # C6680) and p75 (Advanced Targeting Systems, # AB-N07), and negative for iPSC markers Oct4, SSEA3/4, and Sox2 (**Supplement Figure 2 and 3**). When assessing cell characteristics at specific time points of differentiation, cells were collected on specified days by scraping, spun down at 200 × g for 5 minutes, and frozen after aspiration of the supernatant. Successful differentiation of WT and GM3SD variant iPSCs to NCCs was monitored by endpoint analysis for the acquisition of NCC markers and loss of pluripotency markers by immunofluorescence as described below. Marker assessment was performed over multiple independent differentiations and repeated upon thaw of each new aliquot of an iPSC line. In all cases, the iPSCs acquired NCC and lost pluripotency markers as expected based on previous publications [16].

### Immunofluorescence staining

Antibodies and dilutions were as follows: Anti-HNK-1 antibody (Sigma Aldrich, Monoclonal Anti-HNK-1/N-CAM (CD57), # C6680), 1:200 dilution in 5% donkey serum prepared in 0.2% Triton X-100 in PBS (PBS-T); Anti-Sox2 antibody (R&D, # MAB2018), 1:200 dilution in 5% donkey serum in PBS-T; Anti-Oct4 antibody (Santa Cruz, # sc-8628), 1:200 dilution in 5% donkey serum in PBS-T; Anti-Pax6 antibody (Santa Cruz, # sc-81649), 1:200 dilution in 5% donkey serum prepared in 0.05% Tween-20 in PBS (PBS-T); Anti-SSEA3/4 antibody (Santa Cruz, # sc-21704), 1:200 dilution in 5% donkey serum in PBS-T; Anti-p75 antibody (Advanced Targeting Systems, # AB-N07), 1:100 dilution in PBS-T. All antibodies were incubated overnight at 4°C. Immunofluorescent images were captured using an Olympus Fluoview FV1000 laser confocal microscope or a LionheartFX (BioTek) fluorescence microscope.

### Extraction and preparation of GSLs

iPSCs or iPSC-derived NCCs were homogenized and extracted with organic solvents to precipitate proteins and recover GSLs as described previously [18]. Briefly, cell pellets were Dounce homogenized in 50% methanol on ice. Ice-cold water (W), methanol (M) and chloroform (C) were added to give final ratio of C:M:W::4:8:3. The suspension was then transferred into a Teflon-lined screw cap glass tube, agitated for 2 hours at room temperature, and clarified by centrifugation. The supernatant was saved and the resulting pellet was re-extracted three times with fresh 4:8:3 solvent. All supernatants were combined as the lipid extract. Lipid extracts were dried under a nitrogen stream and subjected to saponification to remove glycerophospholipids [19]. Saponification was completed using 0.5M KOH in methanol-water (95:5, M/W, v/v) at 37 °C for 6 hours. Following neutralization with 10% acetic acid on ice, the solution was adjusted to 50% M/W, and then directly loaded onto a Sep-Pak tC18 cartridge column (Waters, Sep-Pak Vac 1cc, 100 mg resin) equipped with a glass syringe (10 ml) that was previously washed with methanol and pre-equilibrated with distilled water. The initial flow-through from the loading was collected and re-applied to the column to enhance recovery. The loaded column was washed with a total of 30 ml of water. GSLs were then eluted with 3 ml of methanol and dried under a nitrogen stream. Free fatty acids resulting from saponification of glycerophospholipids were removed from the dried GSLs by wash with hexane and redrying under nitrogen. Initial analyses of WT and GM3SD GSLs were performed by thin-layer chromatography (TLC) using the following solvent systems: C/M/W (60:35:8) for neutral GSLs; C/M/0.2% CaCl_2_ (55:45:10) for gangliosides; C/M/W (60:40:10) for GSL mixtures. Orcinol–H_2_SO_4_, resorcinol, Dittmer-Lester, and Ninhydrin reagents were used for the detection of sugar, sialic acid, phosphate, and primary amine groups, respectively.

### Analysis of GSLs by mass spectrometry

Nanospray ionization mass spectrometry (NSI-MS) was performed on permethylated glycolipids. For MS of permethylated glycolipids in positive ion mode,∼0.4 nmol of permethylated total glycolipids were dissolved in 50 µl of 1 mM sodium acetate in methanol/water (1:1) for infusion into a linear ion trap mass spectrometer (Orbi-LTQ; ThermoFisher Scientific) using a nanospray source at a syringe flow rate of 0.40 µl/min and capillary temperature set to 210 °C [18–21]. The instrument was tuned with a mixture of permethylated standard neutral or ganglioside GSLs. For fragmentation by collision-induced dissociation (CID) in MS/MS and MS^n^, a normalized collision energy of 30 – 35% was used.

Detection and relative quantification of the prevalence of individual glycolipids were accomplished using the total ion mapping (TIM) and neutral loss scan (NL scan) functionality of the Xcalibur software package version 2.0 (ThermoFisher Scientific) as previously described [18]. For TIM, the m/z range from 600 to 2000 was automatically scanned in successive 2.8 mass unit windows with a window-to-window overlap of 0.8 mass units, which allowed the naturally occurring isotopes of each glycolipid species to be summed into a single response, thereby increasing detection sensitivity. Most glycolipid components were identified as singly, doubly, and triply charged, sodiated species (M + Na) in positive mode. Peaks for all charge states were summed for quantification. Preliminary analysis demonstrated that the major fragment ions in CID MS/MS scans of glycolipid preparations correspond to the neutral loss of the ceramide moiety, leaving intact glycolipid oligosaccharide ions. Therefore, an MS workflow was defined for NL scans in which the highest intensity peak detected by full MS was subjected to CID fragmentation. If an MS/MS profile contained an ion with m/z equivalent to loss of the most prevalent ceramide moiety, MS^n^ fragmentation was initiated. Following this data-dependent acquisition, the workflow returned to full MS, excluded the parent ion just fragmented, and chose the peak of next highest intensity for the same MS/MS and MS^n^ analysis. Graphic representations of GSL monosaccharide residues are consistent with the Symbol Nomenclature For Glycans (SNFG) as adopted by the glycomics and glycobiology communities [22]. Glycomics data and metadata were obtained and are presented in accordance with MIRAGE standards and the Athens Guidelines [23, 24]. All raw mass spectrometric data, both glycomic and proteomic, will be deposited at GlycoPost upon submission for peer-review, Accession #XXXX [25].

### Selective Exo-Enzymatic labeling of cell surface glycoproteins

Selective Exo-Enzymatic Labeling (SEEL) installs a biotin moiety onto the N-linked glycans of cell-surface glycoproteins, allowing their subsequent high-affinity capture and enrichment for proteomic analysis [26, 27]. SEEL and subsequent proteomic analysis of tagged proteins were conducted on WT and p.Glu355Lys iPSCs and NCCs according to previously described methods [27]. Recombinant rat α-(2,6)-sialyltransferase (ST6Gal1) was prepared as previously described [28]. Biotinylated CMP-sialic acid was synthesized as described previously according to [27]. For labeling of cell-surface glycoproteins, approximately 9×10^7^ NCCs from WT and p.Glu355Lys lines were dissociated from the cell culture dish by manual trituration in Dulbecco’s phosphate-buffered saline (DPBS) and the cells were transferred to Eppendorf tubes. The cells were pelleted and resuspended in serum-free DMEM with 42 µg/ml ST6Gal1, 34 µM CMP-Sia-C5-biotin, 13.3 µg/ml BSA, 13.3 µg/ml alkaline phosphatase and 2 µl (in 50% glycerol) *Arthrobacter ureafaciens* (AU) neuraminidase. Each reaction was in a final volume of 350 µl per tube and was incubated for 2 hours at 37 °C. After the SEEL labeling incubation, labeled cells were lysed in RIPA buffer and biotinylated proteins were captured by immunoprecipitation (IP) from 1 mg total lysate protein using anti-biotin antibody. Precipitated proteins were resolved by SDS-PAGE and visualized by silver stain. The resulting gels were cut into two pieces corresponding to regions of the gel that contained proteins with apparent molecular weight between 75-150 kDaltons or greater than 150 kDaltons. Each gel region was then subjected to in-gel trypsinization as previously described [27]. The resulting peptides and glycopeptides from each gel region were separately analyzed by LC-MS/MS using a Lumos Tribrid mass spectrometer (ThermoFisher). Proteins identified by peptides detected at <10 spectral matches (PSMs) were excluded from analysis, as were peptides identified as nuclear or cytosolic by GO annotation (**Supplement Table 1**). Spectral counts for cell surface and secreted proteins that met threshold criteria were normalized to the number of spectral counts detected for human transferrin receptor in WT and p.Glu355Lys IPs.

### Identification of Receptor Tyrosine Kinases in WT and GM3SD cells

Protein was harvested from WT and p.Glu355Lys iPSC and NCC lysates (d19 of differentiation) using the manufacturer’s Receptor Tyrosine Kinase (RTK) array protocol in Lysis buffer 17 (R&D Systems, # ARY001B). RTK arrays were blocked for 1 hour at room temperature in Array Buffer 1 (manufacturer’s protocol). An amount of protein (300 µg) from each cell lysate was diluted in Array Buffer 1 to a volume of 1.5ml and added to the blocked RTK arrays. Arrays were then incubated with gentle rocking overnight at 4 °C. The arrays were then washed 3X with 1X Wash Buffer (manufacturer’s protocol) and subsequently probed with HRP-conjugated pan-phosphotyrosine antibody diluted in Array Buffer 2 (manufacturer’s protocol). Following incubation with gentle rocking for 2 hours at room temperature, the blots were washed with Wash Buffer and RTK capture/phosphorylation was detected by chemiluminescence.

### Western blot analysis

Antibodies for detection of cell surface proteins by western blot of cell lysates were used as provided and were from the following sources: anti-ErbB3 (Cell Signaling, HER3/ErbB3, D22C5 XP Rabbit mAb, # 12708) at 1:1000 dilution with detection by goat anti-mouse secondary at 1:2500 dilution; anti-EGFR (Santa Cruz, EGFR A-10, # sc-373746) at 1:1000 dilution with detection by goat anti-mouse secondary at 1:2500 dilution; anti-caspase 3 (Santa Cruz, caspase-3 Antibody 31A1067, # sc-56053) at 1:1000 dilution with detection by goat anti-mouse secondary at 1:2500 dilution; anti-sortilin (BD Biosciences, Neurotensin Receptor 3 antibody, # 612100) at 1:1000 dilution with detection by goat anti-mouse secondary at 1:2500 dilution; anti-Notch2 (DSHB, Notch2 C651.6DbHN) at 1:500 dilution with detection by goat anti-rat secondary at 1:3000 dilution; anti-β-actin (Santa Cruz, Actin Antibody 2Q1055, # Sc-58673) at 1:5000 dilution with detection by goat anti-mouse secondary at 1:2500 dilution. O-GlcNAcylation of proteins in cell lysates was detected by western blot with anti-O-GlcNAc monoclonal antibody RL-2 (Enzo Life Sciences, ALX-804-11-R100) at 1:1000 dilution with detection by goat anti-mouse secondary at 1:10000. Actin was detected in parallel with O-GlcNAc by incubation with anti-β-actin (Cell Signaling Technology, Actin Antibody 13E5, #4790) at 1:5000 with detection by goat anti-rabbit secondary at 1:10000.

Protein concentrations in lysates were measured by BCA assay and aliquots containing 25 µg were reduced, alkylated, resolved by SDS-PAGE, and transferred to PVDF membranes for probing with primary antibodies. Primary antibody binding was detected by HRP-conjugated secondary antibodies and chemiluminescence. Densitometric quantification was performed with ImageJ Software. For developmental time courses that assessed the change in abundance of receptors across differentiation from iPSC to NCC, signal intensities for receptor proteins were normalized to ß-actin (loading control) and then expressed as log_2_-fold changes relative to the normalized signal intensity detected in iPSC (d0 of differentiation).

### Pharmacologic inhibition of EGFR signaling by erlotinib and of O-GlcNAcase by thiamet-G

WT and p.Glu355Lys iPSCs were differentiated to NCCs for 10 days. Beginning at d10, vehicle (2% DMSO final) or erlotinib in vehicle (100 µM final concentration in media) or thiamet-G in vehicle with or without erlotnib (10 µM thiamet-G ± 100 µM erlotnib final concentration in media) was added to a media change and the cells were incubated for 24 or 48 additional hours prior to harvest for analysis. At harvest timepoints, the media was removed from the wells and centrifuged to collect floating cells. Adherent cells were also scraped and pelleted for collection. Adherent and floating cells were lysed prior to analysis by BCA assay and analyzed separately by western blot for the relative abundance of cleaved and uncleaved caspase 3.

### Statistical analysis

Western blot analyses were performed on cells harvested at indicated time points from between 3-6 independent differentiation courses. Multiple parametric tests were utilized to assess the statistical significance of the resulting data. One-way ANOVA was used to identify significant changes across the three cell populations: WT, p.Glu355Lys, and p.Arg288Ter. Z-scores were calculated and subjected to Z-test to assess the distribution of WT and GM3SD population data for EGFR and ERBB3 expression, where the large number of independent replicates allowed for reasonable estimation of the standard deviation of the entire population. Pairwise T-tests were also used to assess the similarity of one population compared to another for specific features, especially for assessing pharmacologic effects on smaller sample sets. For all three tests, a P-value less than or equal to 0.5 was considered indicative of a significant difference between the tested populations. The results of pairwise T-tests and Z-tests identified the same population differences as significant; for simplicity of presentation, only the pairwise T-tests are presented for direct comparisons. In tables and graphs, population means are reported ± standard error of the mean. For the analysis of EGFR and ERBB3 expression, immunoblot band intensities for each receptor were normalized to actin intensity in order to control for loading variability and then presented as log_2_ of the ratio for each timepoint relative to day 0 of the differentiation course. For the analysis of cleaved Caspase3, the ratio of the immunoblot intensity associated with the cleavage product at 17kD was divided by the ratio of the intensity associated with the intact protein at 34kD detected in the same lane; normalization to actin was not necessary.

## RESULTS

### GM3 synthase deficiency impacts neural-specific GSL biosynthesis

Quantification of the amount and the relative abundance of GSLs in WT and GM3SD cell populations by mass spectrometry demonstrates broad similarity as iPSCs, but highly divergent profiles in the differentiated NCCs population (Table 1, Figure 2). The GSL profiles of iPSCs and NCCs are detectably different by thin-layer chromatography, whether the cells are WT or GM3SD (Figure 3). WT and GM3SD iPSC GSL profiles are dominated by hexosyl-, lactosyl-, globotriaosyl- and globotetraosylceramide (Gb3 and Gb4, respectively). The induced pluripotent stem cell marker SSEA-3, itself a GSL, was detected in WT and GM3SD iPSC GSL profiles. Upon differentiation of WT iPSCs to NCCs, LacCer is efficiently shunted toward the production of GM3, along with the more complex gangliosides derived from GM3, such as GD3, GM1, and GD1 (see Figure 1). Differentiation of GM3SD iPSCs results in accumulation of LacCer without production of GM3. Higher resolution analysis of GSL profiles by NSI-MS^n^ of their permethylated derivatives validates the lack of GM3 in GM3SD iPSCs and NCCs and the resulting lack of a-, b-, or c-series gangliosides (Figure 4,5). The complex sialylated GSLs detected in GM3SD NCCs were detected only as o-series gangliosides, these alternatively sialylated GSLs are sialylated on their external Gal residue (GM1b, GD1c, see Figure 1). In the absence of GM3 and GM3-derived GSLs, the GM3SD NCCs retained high levels of LacCer and elevated levels of Gb3- and Gb4Cer compared to wildtype NCCs.

**Figure 2.**
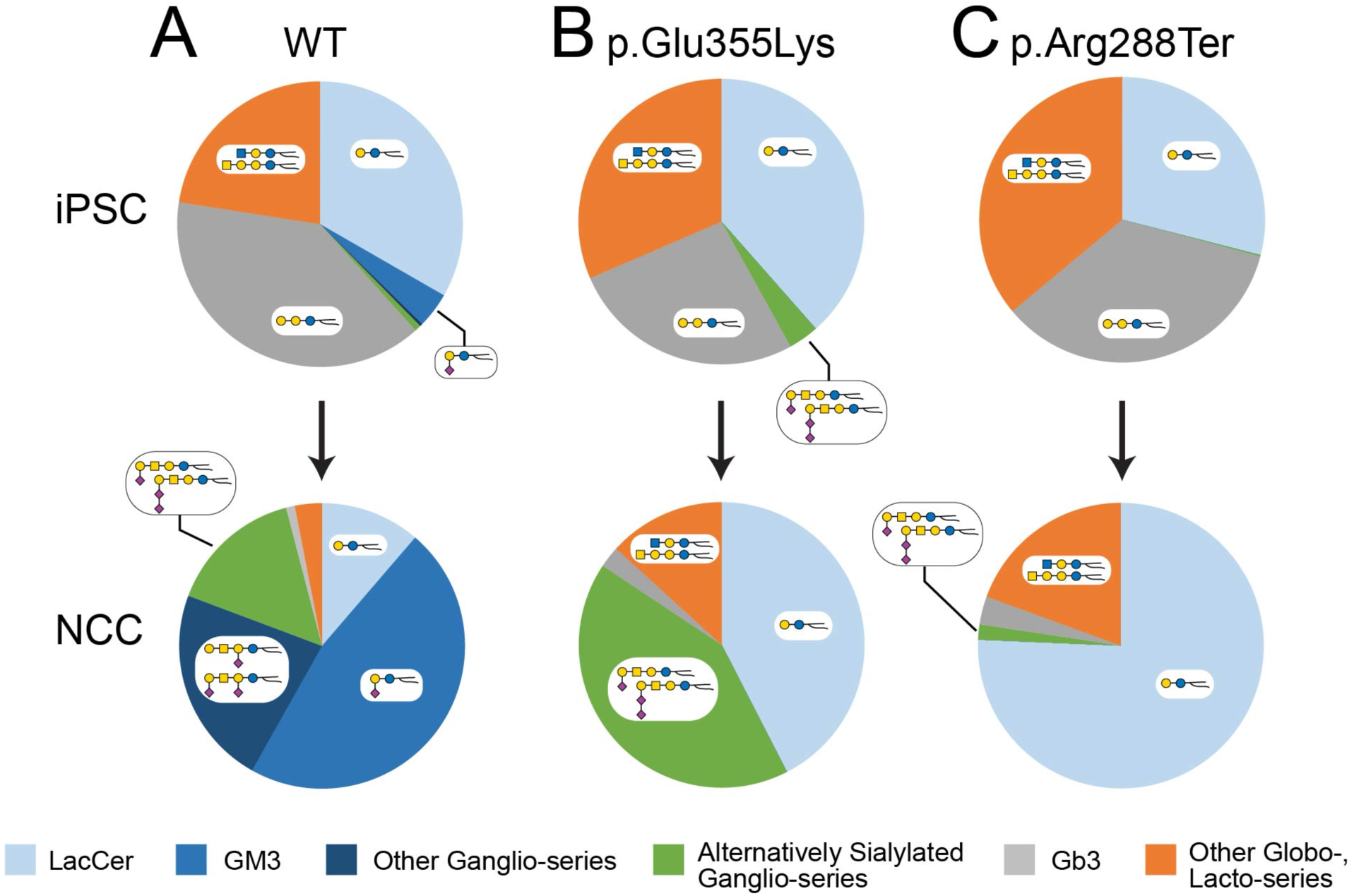
Summary of GSL abundance changes in iPSCs and NCCs of wildtype and GM3SD variants. The relative abundance of the major GSL types detected in iPSCs and NCCs demonstrate relatively minor differences in iPSCs but more significant changes upon differentiation to NCCs. (**A**) As WT iPSCs differentiate to NCCs, decreases in the relative abundance of LacCer, Gb3, and other Globo-, and Lacto-series GSLs are mirrored by increases in GM3 and other Ganglio-series GSLs. (**B,C**) The GSL profiles of GM3SD variant iPSCs are very similar to WT with the exception that GM3 is not detected and p.Glu355Lys cells generate alternatively sialylated Ganglio-series GSLs that are independent of the activity of ST3GAL5. Differentiation of GM3SD variant iPSCs to NCCs is accompanied by maintenance of LacCer as a major GSL and by complete lack of Ganglio-series GSLs that depend on ST3GAL5 activity. The p.Glu355Lys NCCs are more efficient at generating alternatively sialylated GSLs (do not require ST3GAL5), while p.Arg288Ter NCCs accumulate LacCer without diversion to other biosynthetic pathways.

**Figure 3.**
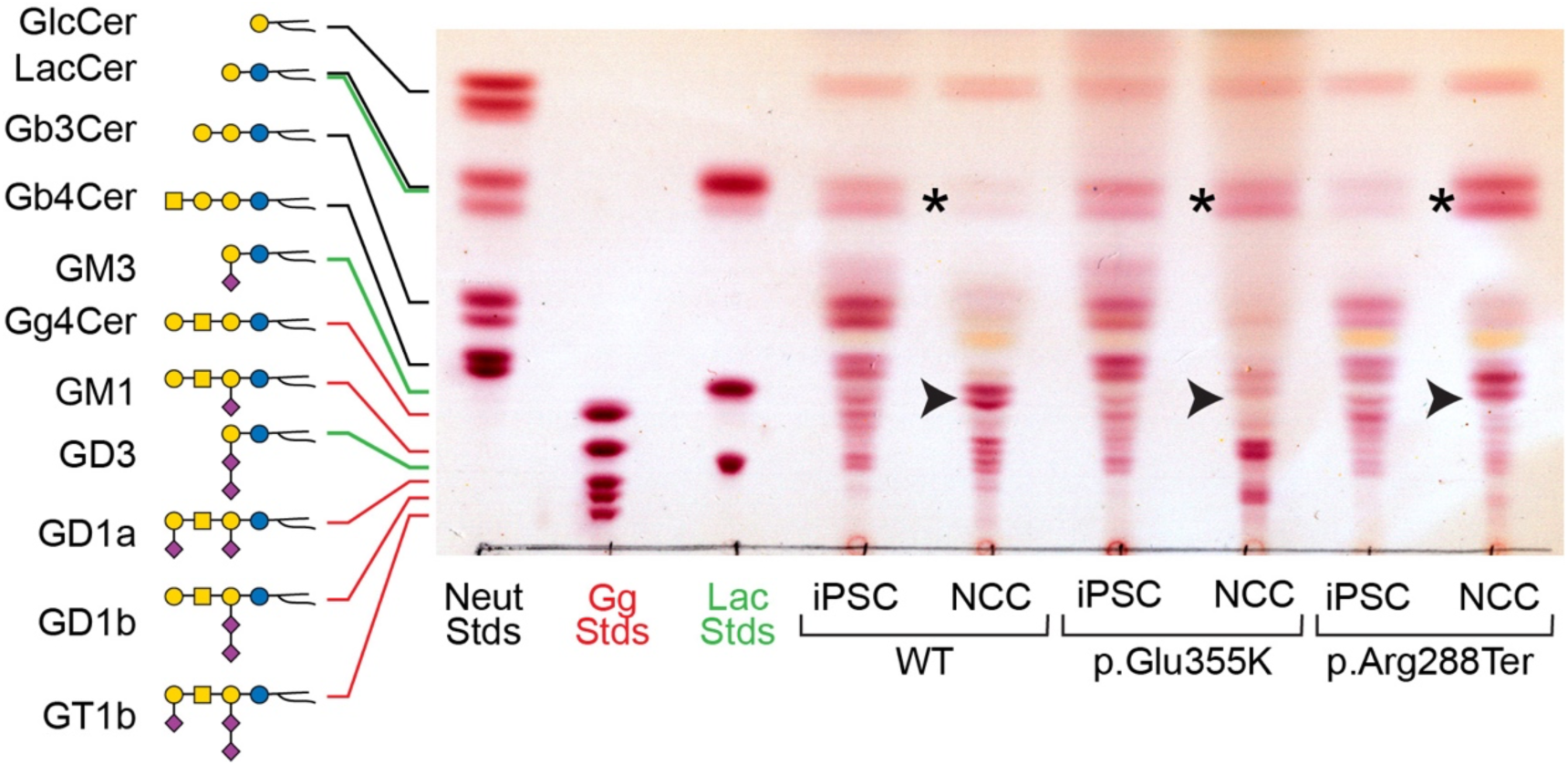
iPSCs and NCCs derived from GM3SD variants fail to produce GM3 and accumulate LacCer. GSLs extracted from WT and GM3SD patient cell types were resolved by thin-layer chromatography and visualized by staining with orcinol reagent. The mobility of three sets of GSL standards are shown for reference (***Neut***, GSLs with neutral cores of increasing length up to globotetraosylceramide; ***Gg***, GSLs with gangliotetraose cores carrying 0, 1, 2, or 3 sialic acids; ***Lac***, GSLs with a LacCer core carrying 0, 1, or 2 sialic acids). In iPSCs of all three genotypes, the GSL profile is dominated by LacCer and globo-series GSLs. Upon differentiation of WT iPSCs to NCCs, LacCer abundance decreases (***asterisk***) and the appearance of gangliosides is evident, among which GM3 is the most abundant (***arrowhead***). NCCs derived from both GM3SD genotypes retain high levels of LacCer and do not generate GM3.

**Figure 4.**
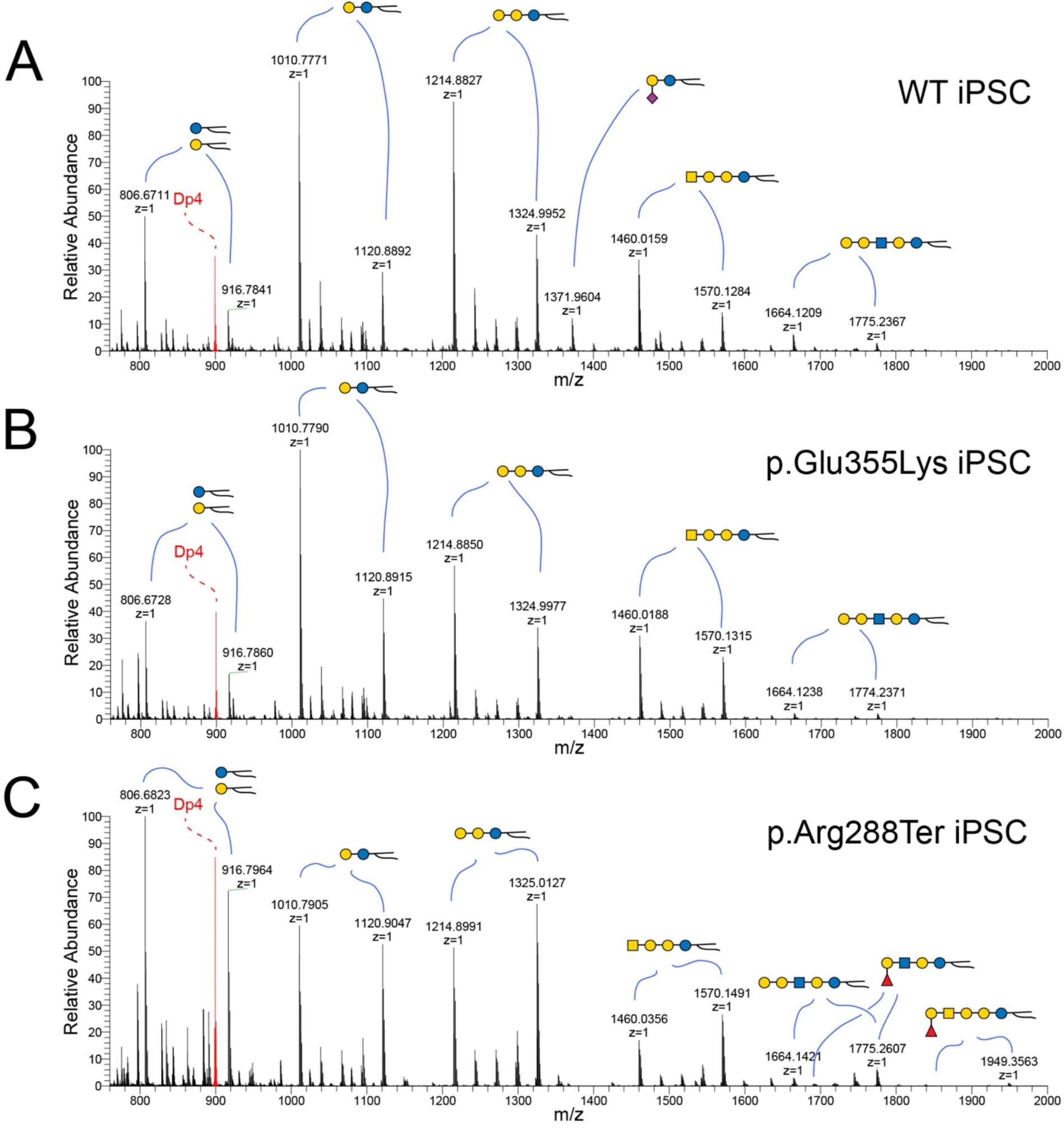
NSI-MS analysis of GSLs extracted from iPSCs detects minor differences in GSL biosynthesis and ceramide compositions. Mass spectra were obtained for permethylated intact GSLs extracted from WT (**A**), p.Glu355Lys (**B**), and p.Arg288Ter (**C**) iPSCs. GSLs with the same glycan headgroup are detected as families of related structures with differing ceramides (***highlighted by blue lines***). GM3 is a minor component of the GSL profile of WT iPSCs (***m/z 1372 in A***) but is not detected in either of the GM3SD iPSCs. Both of the GM3SD iPSCs produce a higher relative abundance of longer ceramides; compare the ratio of the LacCer form at m/z 1011 to the form at m/z 1121 across all three iPSCs. The shift in ceramide to higher mass is more pronounced in p.Arg288Ter than in p.Glu355Lys.

**Figure 5.**
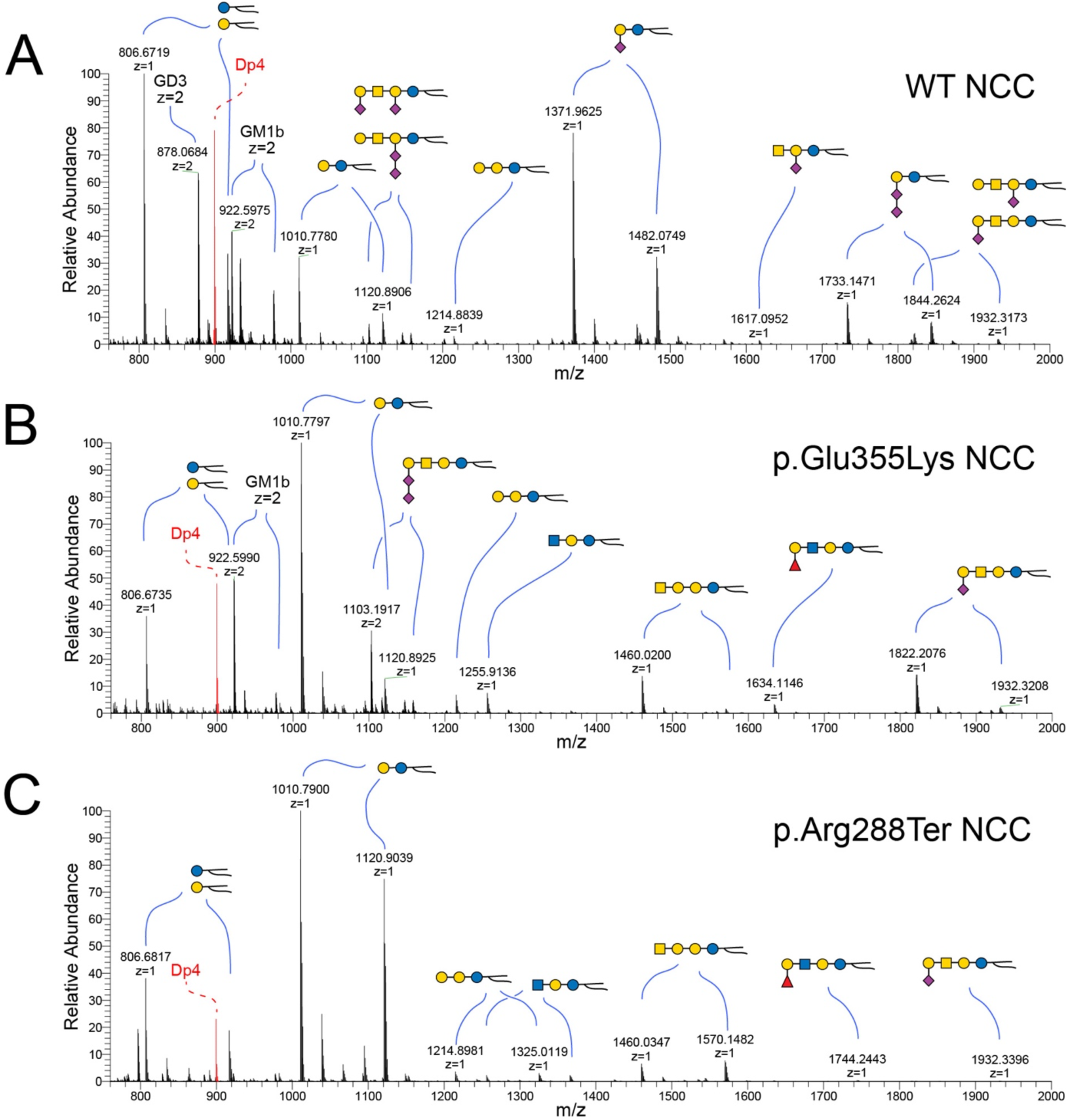
NSI-MS analysis of GSLs extracted from NCCs highlights neural cell-specific alterations in GSL biosynthesis and ceramide compositions. Mass spectra were obtained for permethylated intact GSLs extracted from WT (**A**), p.Glu355Lys (**B**), and p.Arg288Ter (**C**) NCCs. Upon differentiation of WT iPSCs to NCCs, GM3 becomes the most abundant GSL, but is not detectable in GM3SD NCCs. Both populations of GM3SD NCCs retain elevated LacCer (***m/z 1011-1103***). p.Glu355Lys NCCs synthesize alternatively sialylated GSLs, such as GM1b (***m/z 1822-1932***) and GD1c (***m/z 1103-1159, z=2***), at higher levels than p.Arg288Ter while p.Arg288Ter retains higher mass ceramides to a greater extent than p.Glu355Lys.

**Table 1.** GSL abundance in iPSCs and NCCs of wildtype and GM3SD variant cells.

| GSL<sup>a</sup> | Cell type and Genotype |  |  |  |  |  |  |  |  |  |  |  |
| --- | --- | --- | --- | --- | --- | --- | --- | --- | --- | --- | --- | --- |
|  | iPSC |  |  |  |  |  | NCC |  |  |  |  |  |
|  | Wildtype |  | p.Glu355Lys |  | p.Arg288Ter |  | Wildtype |  | p.Glu355Lys |  | p.Arg288Ter |  |
|  | %<sup>b</sup> | pmol<sup>c</sup> | % | pmol | % | pmol | % | pmol | % | pmol | % | pmol |
| **LacCer** | 33.3 | 277.8 | 38.5 | 230.4 | 29.0 | 36.1 | 11.3 | 35.1 | 42.4 | 138.4 | 75.7 | 175.7 |
| **Gb3** | 39.1 | 326.1 | 26.6 | 158.9 | 34.8 | 43.3 | 1.0 | 3.1 | 2.4 | 7.9 | 3.2 | 7.4 |
| **Lc3** | --<sup>d</sup> | -- | -- | -- | 0.2 | 0.2 | 0.4 | 1.2 | 2.8 | 9.0 | 2.1 | 4.8 |
| **Gb4, Lc4** | 20.8 | 173.8 | 31.5 | 188.4 | 21.2 | 26.4 | 2.7 | 8.3 | 7.8 | 25.6 | 16.0 | 37.1 |
| **Fuc-Lc4** | 1.7 | 14.1 | -- | -- | 14.7 | 18.3 | -- | -- | 2.6 | 8.5 | 1.4 | 3.2 |
| **GM3** | 4.0 | 33.6 | -- | -- | -- | -- | 46.9 | 145.6 | -- | -- | -- | -- |
| **GM2** | -- | -- | -- | -- | -- | -- | 0.3 | 1.0 | -- | -- | -- | -- |
| **GM1a** | 0.3 | 2.6 | -- | -- | -- | -- | 1.5 | 4.6 | -- | -- | -- | -- |
| **GM1b** | 0.7 | 6.0 | 3.4 | 20.6 | 0.2 | 0.2 | 13.4 | 41.8 | 28.2 | 92.1 | 1.6 | 3.7 |
| **GD1a** | -- | -- | -- | -- | -- | -- | 20.5 | 63.7 | -- | -- | -- | -- |
| **GD1b** | -- | -- | -- | -- | -- | -- | 0.3 | 0.8 | -- | -- | -- | -- |
| **GD1c** | -- | -- | -- | -- | -- | -- | 1.8 | 5.5 | 13.6 | 44.5 | 0.1 | 0.3 |

<sup>a</sup>Graphical representations of each quantified glycosphingolipid are shown in Figure 1, along with the abbreviation used in this table, except for Fuc-Lc4 which denotes a terminally fucosylated lactotetraosylceramide (see Figures 3, 4).
<sup>b</sup>GSL relative abundance, denoted “%,” is given as the percent that each individual GSL contributes to the total lipid profile for each cell type/genotype.
<sup>c</sup>GSL amount, denoted “pmol,” is given as pmol/1x10<sup>6</sup> cells.
<sup>d</sup>Double dash denotes GSL not detected.

GSL ceramide moieties impart physicochemical characteristics that facilitate their incorporation into specialized membrane microdomains referred to as lipid rafts and as detergent resistant membranes [29]. Heterogeneity in GSL ceramides derives from the chemical nature of the sphingosine base and its amide-linked fatty acid. These components can differ based on degree of saturation, hydrocarbon chain length, and presence or absence of a sphingosine hydroxyl group. The heterogeneity of GSL ceramides is well resolved by NSI-MS^n^, in which each GSL glycan head group is detected in association with a family of m/z values that reflect lipid differences (Figure 4,5). The ratio of longer chain fatty acid-containing ceramides to shorter chain-containing ceramides was higher in the iPSCs of GM3SD patients compared to wild-type; LacCer and globoside Gb4 exhibit this difference particularly well in iPSCs (Figure 4). For both iPSCs and NCCs, p.Arg288Ter had a higher abundance of longer chain fatty acid-containing ceramides than p.Glu355Lys, with the ceramide distribution of p.Glu355Lys NCCs closely approximating the ceramide distribution of WT (Figure 5).

While the GSL profiles of p.Glu355Lys and p.Arg288Ter iPSCs and NCCs presented similar deficiencies in GM3 and the major GM3-derived GSLs, more subtle differences were detected between the cells derived from the two GM3SD variants. The alternatively sialylated gangliosides GM1b and GD1c were increased in p.Glu355Lys NCCs compared to p.Arg288Ter NCCs (Table 1, Figure 2). Additionally, GSLs with N-acetyl-lactosamine (LacNAc) reapeats uncapped by sialic acid at their non-reducing termini were more abundant in p.Arg288Ter NCCs, whereas extended LacNAc structures were more likely to be sialylated in p.Glu355Lys NCCs (Figure 6, see **Supplement Figure 4** for validation of the extended LacNAc GSL structures). In general, the major shifts in GSL profiles were conserved across the two GM3SD variants when compared to WT, but p.Glu355Lys NCCs synthesized a higher abundance of alternatively sialylated gangliosides and p.Arg288Ter cells retained greater relative abundance of LacCer and ceramides with higher mass.

**Figure 6.**
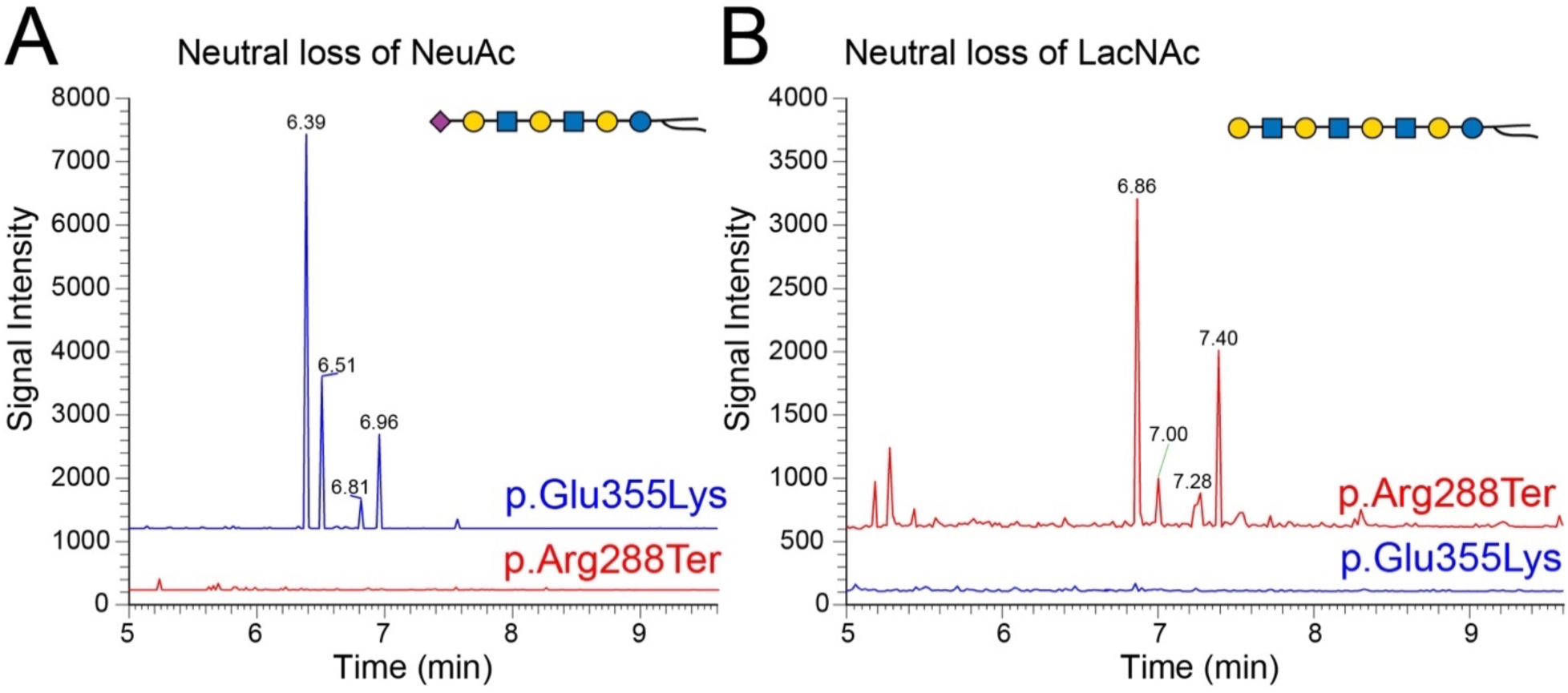
Unique GSL changes in GM3SD Variants. The results of Total Ion Monitoring workflows (TIM-chromatograms) were filtered for neutral loss of NeuAc or LacNAc. (**A**) Filtering for loss of NeuAc demonstrates that GSLs with extended LacNAc repeats capped with sialic acid are detected at higher abundance in p.Glu355Lys NCCs than in p.Arg288Ter NCCs. (**B**) Filtering for loss of LacNAc disaccharide demonstrates that non-sialylated forms of GSLs with LacNAc repeats are more abundant in p.Arg288Ter than in p.Glu355Lys NCCs. MS2 fragmentation profiles support the structural assignments at the inidicated TIM detection times (***Supplement Figure 4***). These less abundant unique GSLs, like the more abundant GSLs in NCCs, also show greater enrichment of higher mass ceramides in p.Arg288Ter than in p.Glu355Lys (e.g., compare the ratio of peaks at 6.96/6.39 in **A** to 7.40/6.86 in **B**).

### GM3 synthase deficiency alters the cell surface proteome

GM3 and several of its downstream GSL products are components of lipid rafts, plasma membrane microdomains enriched in proteins, frequently GPI-anchored, that participate in multiple cellular functions including adhesion and signaling. We employed a robust cell-surface capture technology (Selective Exo-Enzymatic Labeling or SEEL) to assess whether altered GSL profiles in GM3SD cells impacted the abundance of cell surface proteins, in particular those associated with lipid rafts [29–32]. SEEL installs a biotin moiety onto desialylated N-linked glycans of cell surface glycoproteins, allowing their subsequent high-affinity capture and enrichment for proteomic analysis [26, 27]. We applied SEEL to NCCs derived from WT and p.Glu355Lys iPSCs at day 35 of differentiation, captured the resulting biotinylated proteins by immunoprecipitation with anti-biotin antibody, and resolved the precipitated proteins by SDS-PAGE (Figure 7A).

**Figure 7.**
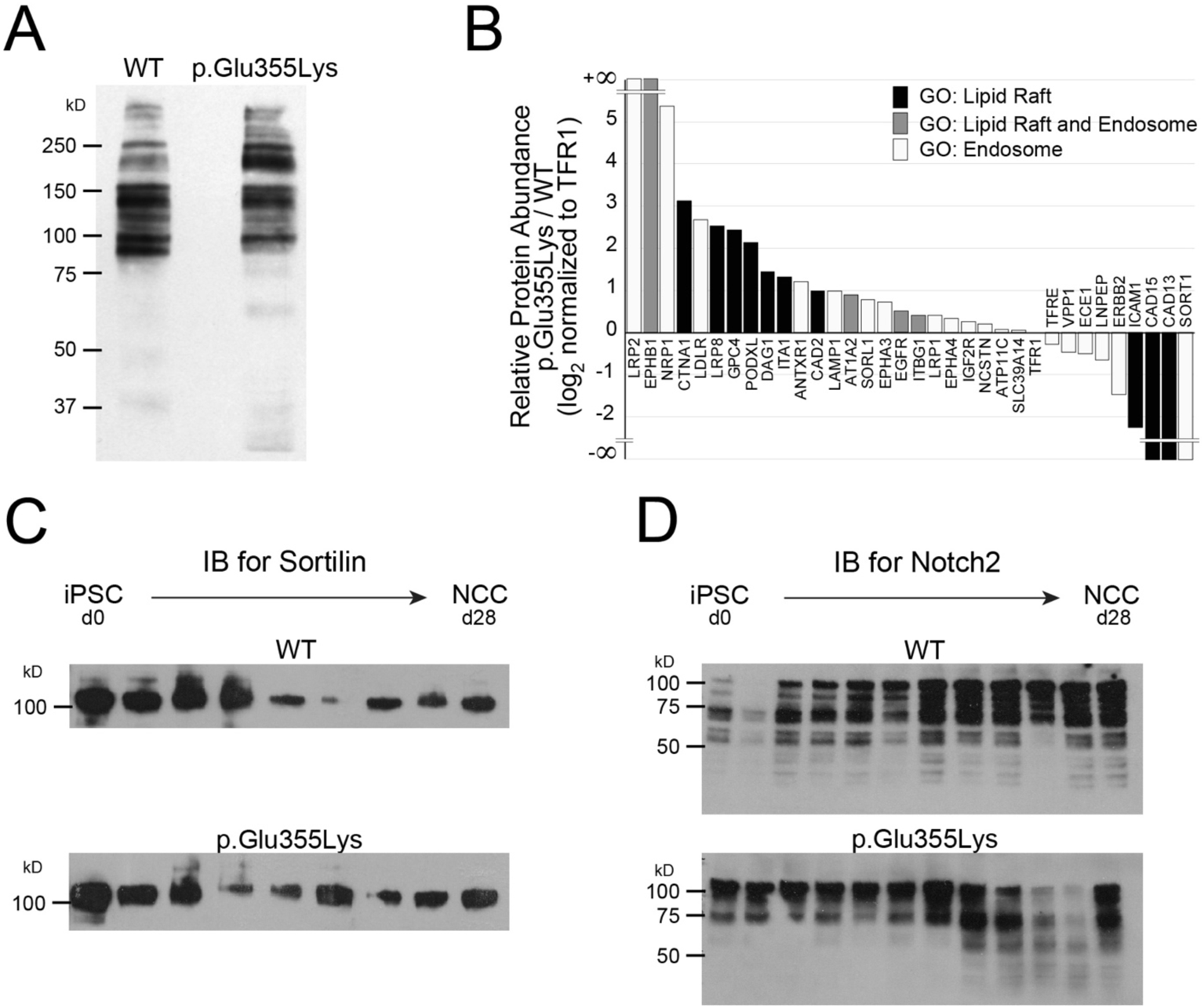
Cell surface protein abundance is altered in GM3SD cells. NC cells differentiated from WT or p.Glu355Lys iPSCs were subjected to selective-exoenzymatic labeling (SEEL) to install a biotin moiety specifically on cell surface glycoproteins. (**A**) Labeled proteins were resolved by SDS-PAGE and probed by western blot with anti-biotin antibody, revealing the presence of changes in the abundance of membrane proteins. (**B**) Gel-resolved proteins were harvested by in-gel digestion with trypsin and identified by LC-MS/MS. Identified proteins assigned to the indicated Gene Ontology (GO) categories were compared to assess the integrity of lipid rafts and the endosomal pathway. (C) One of the proteins that was not detected by cell surface labelling of p.Glu355Lys cells, SORT1, was detectable in whole cell lysates. (D) Notch2, a cell surface receptor important for NC cell differentiation, was also undetectable by surface labeling of p.Glu355Lys cells but found in whole cell lysates (***Supplement Table 1***).

Subsequent in-gel tryptic digestion and LC-MS/MS analysis identified 120 proteins in WT and 160 proteins in p.Glu355Lys detected at > 10 spectral counts (range 10-460). Gene Ontology (GO) analysis indicated that 96% of the identified proteins were cell surface or cell-surface associated and 4% were nuclear or cytoplasmic. Thus, our SEEL method achieved a stringent enrichment of cell-surface glycoproteins and provided a pool of protein identities for further analysis (**Supplement Table 1**). After filtering the total protein identifications to exclude the low-level of contamination with cytoplasmic and nuclear proteins, quantification by spectral counts demonstrated that 73% of the identified cell surface proteins were increased and 27% were decreased in abundance in p.Glu355Lys NCCs compared to WT NCCs.

To assess the impact of GM3SD on the membrane raft and the endocytic pathway proteome, we filtered the identified membrane proteins for GO assignments to the ontologic terms “membrane raft” (GO:0045121) and “endosome” (GO:0005768). Of the total protein identifications, 28% of WT (33 proteins) and 20% of p.Glu355Lys (32 proteins) were annotated with these GO classifications. Among the lipid raft protein identifications, 11 were increased and 3 were decreased in p.Glu355Lys compared to WT (Figure 7B). Among the raft-associated proteins, several adhesion receptors (CAD13, CAD15, ICAM1) were detected in WT NCCs that were absent or significantly reduced in p.Glu355Lys NCCs, while other adhesion receptors (DAG1, ITA1, CAD2) were enriched in p.Glu355Lys NCCs compared to WT. Several endocytic receptors, signaling, and adaptor proteins were increased in abundance at the plasma membrane in p.Glu355Lys NCCs in comparison to WT, while a single signaling receptor, a receptor tyrosine kinase (ERBB2) and two peptide hormone processing enzymes (ECE1, LNPEP) were reduced.

A major component of raft/endosomal membrane trafficking, sortilin (SORT1), was not detected on p.Glu355Lys NCCs following SEEL-based enrichment. However, western blot analysis of whole cell lysates demonstrated that p.Glu355Lys cells express SORT1 at levels similar to WT NCCs (Figure 7C). Likewise, Notch2, a cell surface receptor important for NCC differentiation into peripheral neurons and other NC-derived cell types, was also not detected following SEEL in E335K NCC cells, despite being expressed and detected in whole cell lysates (Figure 7D). These results indicate that the loss of cell surface residence for these receptors in GM3SD cells was not due to lack of expression, but rather to decreased stability of these proteins at a plasma membrane of altered GSL composition.

### GM3 synthase deficiency disrupts protein O-GlcNAcylation, a downstream effector of multiple cell signaling pathways

Protein O-GlcNAcylation is a dynamic modification found on many nuclear and cytoplasmic proteins and is responsive to a broad range of cellular signaling activities, similar to protein phosphorylation. To assess the fidelity of protein O-GlcNAcylation in the absence of complex gangliosides, we probed whole cell lysates for the presence of O-GlcNAc in iPSCs and NCCs derived from WT and GM3SD cells (Figure 8A). For both WT and GM3SD cells, the abundance of O-GlcNAc modified proteins increased upon differentiation of iPSCs to NCCS (Figure 8A,B). The fold-increase in O-GlcNAcylation comparing NCCs to iPSCs was comparable for WT, p.Glu355Lys, and p.Arg288Ter: 3.4 ± 0.9 in WT, 4.1 ± 1.1 in p.Glu355Lys, and 3.6 ± 0.8 in p.Arg288Ter cells (p>0.05 for pairwise comparisons, n=3 independent differentiations for each cell type). While the fold-increases where similar, the total amount of O-GlcNAc modification detected at the end of each separate differentiation course was markedly different, although the population differences did not reach statistical significance (Figure 8B). The apparent molecular weight profile of proteins detectable by western blot of one-dimensional SDS-PAGE gels was different in iPSCs and NCCs, but was not qualitatively altered across genotypes.

**Figure 8.**
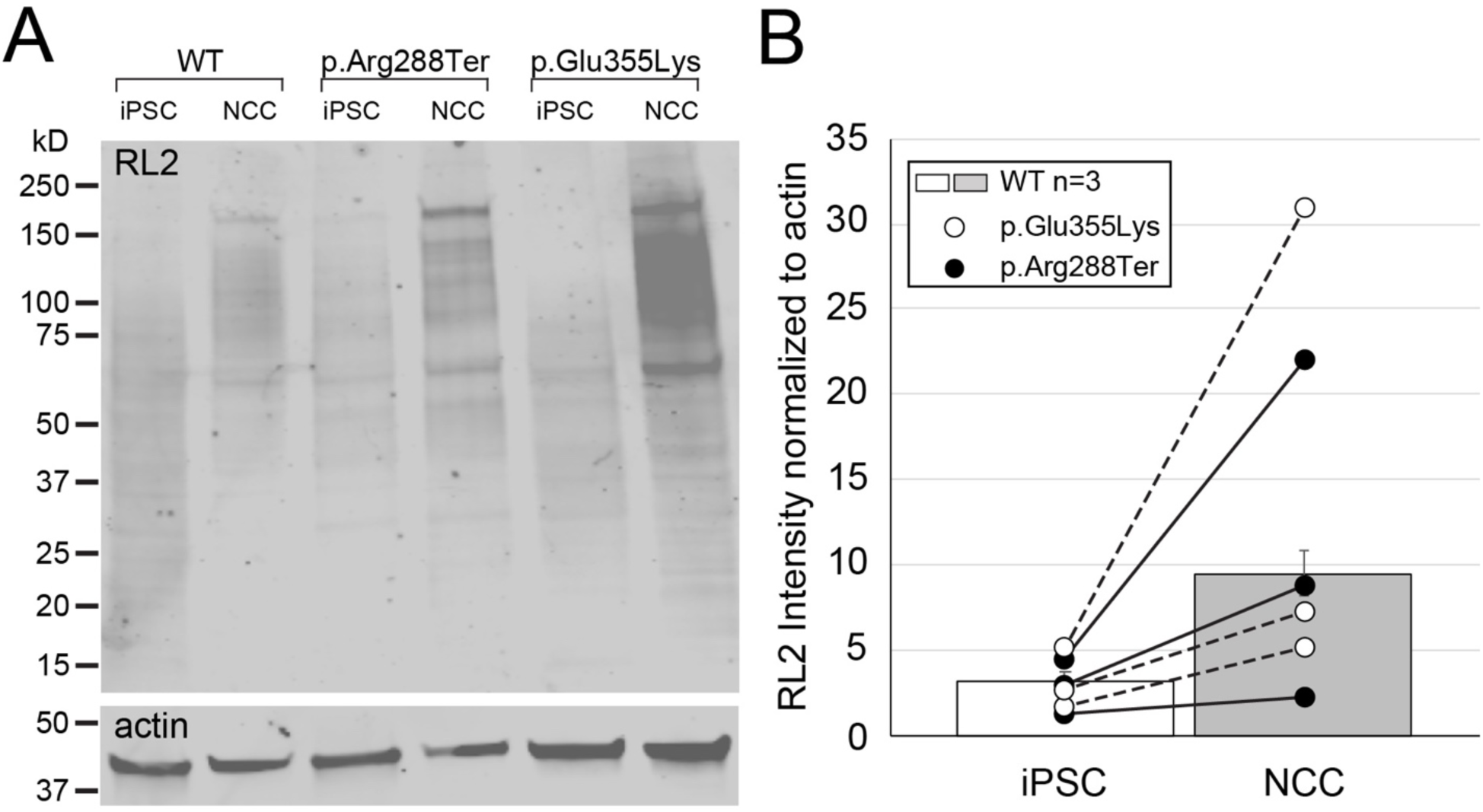
Protein O-GlcNAcylation increases upon differentiation of iPSCs to NCCs. (**A**) Whole cell lysates from WT and GM3SD cells were resolved by SDS-PAGE, blotted, and probed with RL2, a monoclonal antibody that recognizes proteins modified by O-GlcNAc. As iPSCs differentiate to NCCs, the abundance of O-GlcNAcylated proteins increases and the profile of modified proteins also shifts to higher molecular weight. (**B**) As in WT cells, O-GlcNAc levels in GM3SD cells also increase upon differentiation to NC, but the absolute amount of O-GlcNAc is more variable. Individual differentiation experiments for GM3SD cells are shown connected by a dashed or solid line (n=3 independent differentiations). GM3SD cell values are plotted on top of bars that represent the mean ± SEM for WT cells (n=3 independent differentiations).

### Receptor tyrosine kinases are impacted by GM3 synthase deficiency

To assess whether GM3SD broadly impacts cell surface signaling events that lie upstream of protein O-GlcNAcylation, we used an array-based approach to query receptor tyrosine kinase (RTK) activity in whole-cell lysates prepared from WT and p.Glu355Lys iPSCs and NCCs at day 19 of differentiation. This method takes advantage of immobilized antibodies to several different RTKs and detection of the phosphorylation status of these captured RTKs following incubation of the blots with a pan-phosphotyrosine antibody. The major RTKs detected in WT and GM3SD iPSC lysates were EGFR, ERBB3, INSR, and IGF2, each of which were detected at approximately equal intensity comparing WT to GM3SD (Figure 9A). Upon differentiation to NCCs, ERBB3 intensity was increased, while EGFR, INSR, and IGF1 intensities were not substantially modified (Figure 9B). EGFR was similar to WT in GM3SD NCC lysates, while INSR and IGF-1 intensities were slightly decreased. Most striking, however, was the loss of ERBB3 detection in GM3SD NCC lysates. This array-based method reports changes in the overall phosphorylation of the represented RTKs but does not resolve whether the changes relate to altered activity or differences in protein expression. Therefore, we independently assessed the steady-state levels of EGFR and ERBB3 receptors during NCC differentiation.

**Figure 9.**
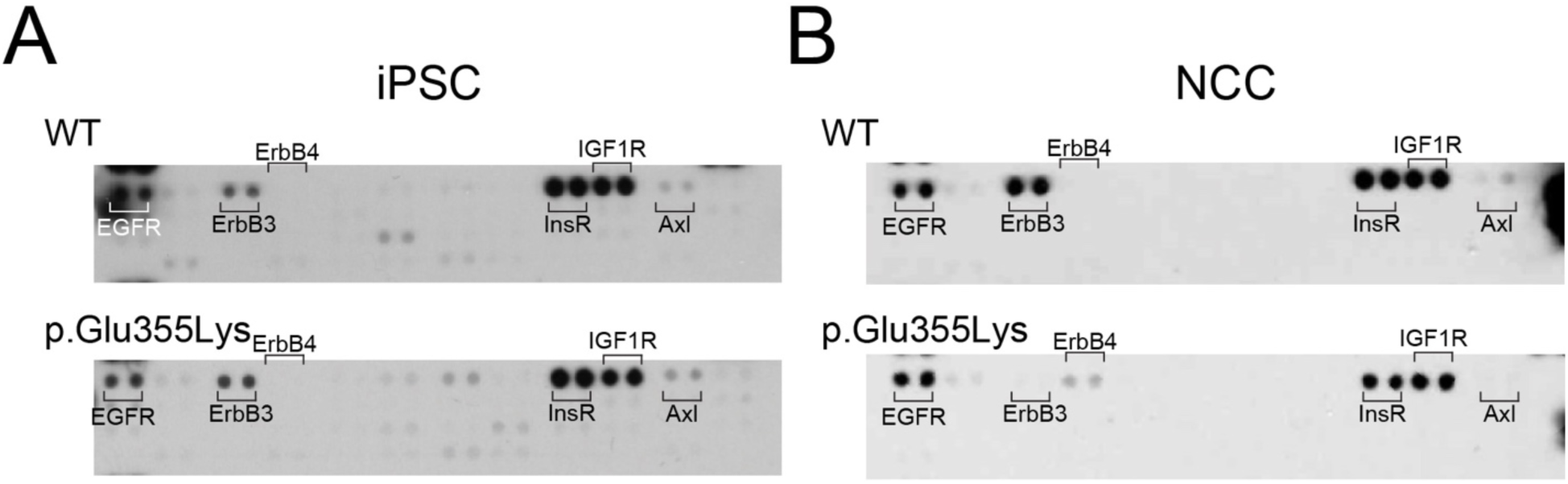
Expression/phosphorylation of receptor tyrosine kinases in WT and GM3SD cells. Blots bearing immobilized anti-RTK antibodies were overlayed with whole cell lysates from (**A**) iPSCs or (**B**) NCCs harvested at d19 of differentiation. Bound RTKs were then detected by sandwich with anti-phosphotyrosine antibody. Signal associated with ERBB3 increases in WT cells upon differentiation to NCCs, but ERBB3 is reduced in p.Glu355Lys NCCs. Other more subtle changes in RTK detection were also evident in p.Glu355Lys NCCs compared to WT.

### GM3 synthase deficiency uncouples EGFR and ERBB3 from neural crest differentiation

EGFR (also known as ERBB1) possesses a binding domain in its extracellular region that interacts with gangliosides, in particular GM3 [33]. EGFR also forms a heterodimer with ERBB3, thought to be a poor kinase by itself, to facilitate signaling in response to heregulin/neuregulin binding [34–36]. To understand whether the differences in ERBB3 detected on the array blots resulted from altered phosphorylation or from differential protein expression/stability, we probed cell lysates with phosphorylation-independent antibodies for EGFR and ERBB3 across the time course of differentiation from iPSCs to NCCs (Figure 10). As WT iPSCs differentiate toward NCCs, they exhibit a reproducible and complementary expression profile in which EGFR increases early in differentiation compared to iPSCs and then decreases upon completion of the differentiation process (Figure 10A **and** 10B). ERBB3 mirrors the EGFR profile, exhibiting an early decrease in abundance that transforms into increased abundance in NCCs compared to iPSCs. The evolving time course of EGFR and ErbB3 expression during differentiation from iPSCs to NCCs was highly reproducible across biological replicates, with the maximum variability occurring during times of greatest change in receptor expression (d0-d15 of differentiation).

**Figure 10.**
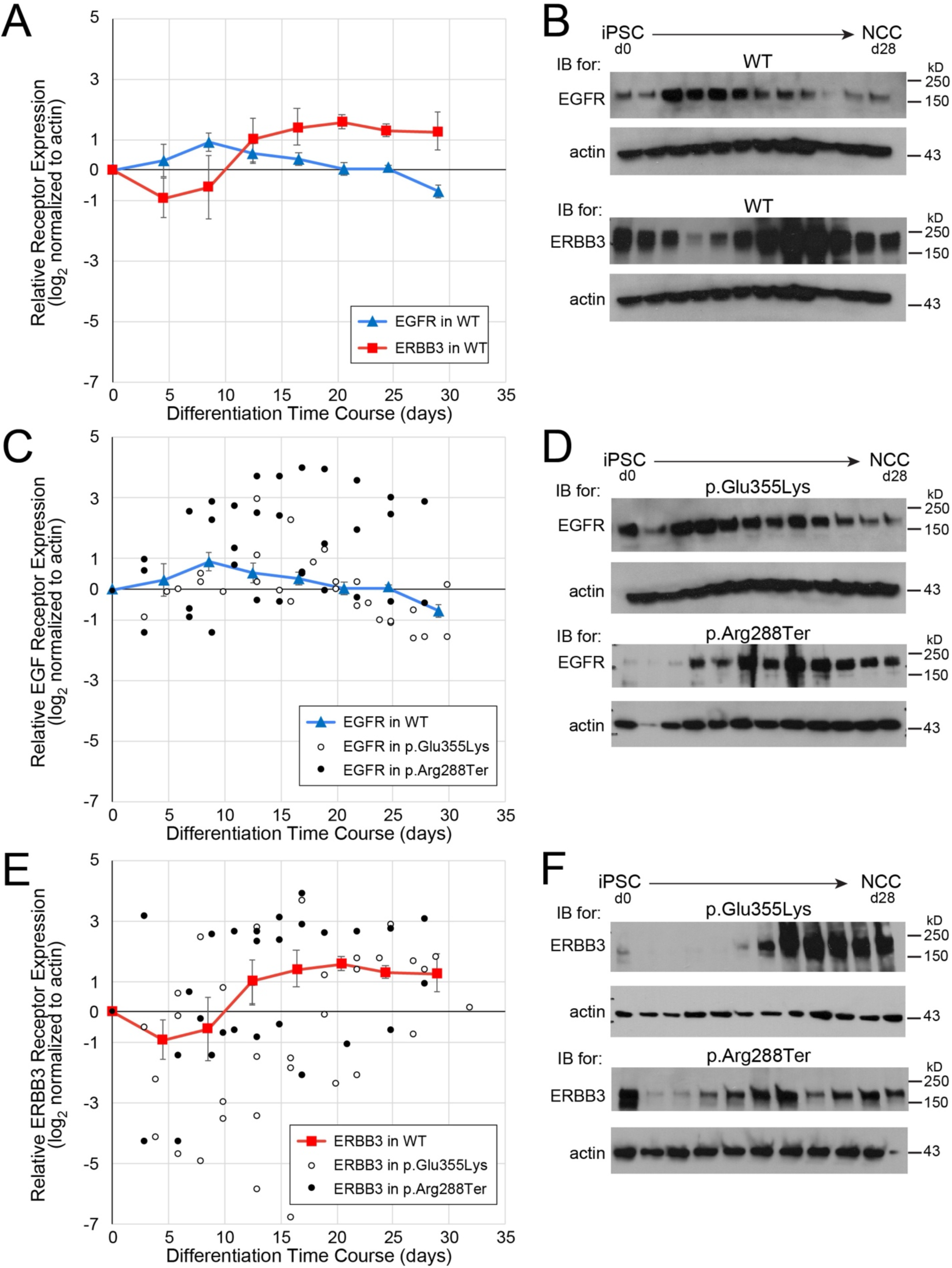
Dynamics of EGFR and ERBB3 expression during neural crest differentiation. Protein-specific, not phospho-specific, antibodies against EGFR and ERBB3 were used to quantify receptor expression in WT and GM3SD cells across full time courses of differentiation from iPSCs to NCCs (n=3 independent differentiations for each cell type). (**A**) EGFR and ERBB3 expression mirror each other as WT iPSCs differentiate to NCCs. For simplicity of presentation and to provide a baseline reference for comparison with GM3SD cells, WT values were combined into bins that span 4 days of differentiation (mean ± SEM, n=4-6 determinations for each bin, total of 30 data points for each receptor). (**B**) Representative EGFR and ERBB3 western blots from one differentiation course of WT cells. (**C**) Individual time points for EGFR abundance are plotted for three independent differentiations of GM3SD cells (n=27 data points for p.Glu355Lys, n=32 data points for p.Arg288Ter); WT EGFR profile is reproduced from panel A for reference. (**D**) Representative EGFR western blots from one differentiation course of p.Glu355Lys and p.Arg288Ter cells. (**E**) Individual time points for ERBB3 abundance are plotted for three independent differentiations of GM3SD cells (n=37 data points for p.Glu355Lys, n=32 data points for p.Arg228Ter); WT ERBB3 profile is reproduced from panel A for reference. (**F**) Representative ERBB3 western blots from one differentiation course of p.Glu355Lys and Arg288Ter cells.

We assessed EGFR and ERBB3 expression in p.Glu355Lys and p.Arg288Ter cells across the same differentiation course from iPSCs to NCCs and compared the GM3SD cells to WT (Figure 10C-F **and** Table 2). Both the p.Glu355Lys and p.Arg288Ter cell populations exhibit significant variation from WT across all timepoints of differentiation. For EGFR, significant differences across the three cell populations were detected during mid (d11-22) and late (d3-32) stages of differentiation (ANOVA p<0.01 and p<0.001, respectively). Pairwise comparisons demonstrated that EGFR expression in p.Glu355Lys was significantly decreased compared to WT at late stages (p<0.05), while expression in p.Arg288Ter was significantly increased in mid and late stages (p<0.01 and p<0.05, respectively). For ERBB3, significant differences were also detected in GM3SD cells compared to WT, especially at mid stages of differentiation (ANOVA p<0.05). Pairwise comparisions demonstrated that ERBB3 expression in p.Glu355Lys was significantly reduced at this stage (p<0.05) but p.Arg288Ter was not significantly different from WT. Pairwise comparisons of EGFR and ERBB3 expression between p.Glu355Lys and p.Arg288Ter further highlight subtle differences between the two GM3SD cell populations, with EGFR expression more highly impacted in p.Arg288Ter and ERBB3 more impacted in p.Glu355Lys cells. Furthermore, the deviation detected for EGFR and ERBB3 in the p.Glu355Lys cells is consistent with the RTK blot, namely ERBB3 was affected more than EGFR at d19 of differentiation (see Figure 9).

**Table 2.**
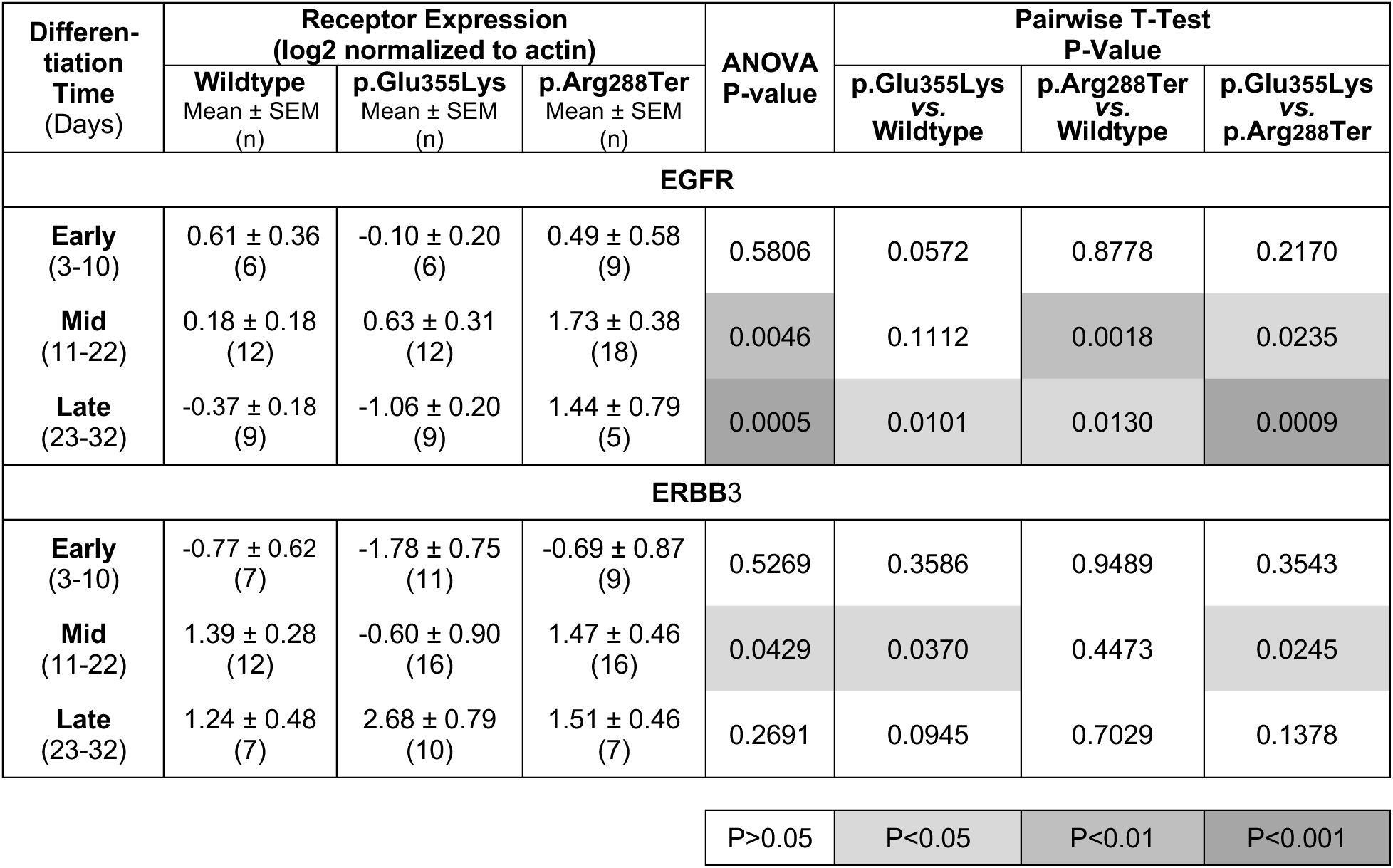
EGFR and ERBB3 expression in iPSCs and NCCs of wildtype and GM3SD variant cells.

| Differentiation Time (Days) | Receptor Expression (log2 normalized to actin) |  |  | ANOVA P-value | Pairwise T-Test P-Value |  |  |
| --- | --- | --- | --- | --- | --- | --- | --- |
|  | Wildtype Mean ± SEM (n) | p.Glu355Lys Mean ± SEM (n) | p.Arg288Ter Mean ± SEM (n) |  | p.Glu355Lys vs. Wildtype | p.Arg288Ter vs. Wildtype | p.Glu355Lys vs. p.Arg288Ter |
| EGFR |  |  |  |  |  |  |  |
| Early (3-10) | 0.61 ± 0.36 (6) | -0.10 ± 0.20 (6) | 0.49 ± 0.58 (9) | 0.5806 | 0.0572 | 0.8778 | 0.2170 |
| Mid (11-22) | 0.18 ± 0.18 (12) | 0.63 ± 0.31 (12) | 1.73 ± 0.38 (18) | 0.0046 | 0.1112 | 0.0018 | 0.0235 |
| Late (23-32) | -0.37 ± 0.18 (9) | -1.06 ± 0.20 (9) | 1.44 ± 0.79 (5) | 0.0005 | 0.0101 | 0.0130 | 0.0009 |
| ERBB3 |  |  |  |  |  |  |  |
| Early (3-10) | -0.77 ± 0.62 (7) | -1.78 ± 0.75 (11) | -0.69 ± 0.87 (9) | 0.5269 | 0.3586 | 0.9489 | 0.3543 |
| Mid (11-22) | 1.39 ± 0.28 (12) | -0.60 ± 0.90 (16) | 1.47 ± 0.46 (16) | 0.0429 | 0.0370 | 0.4473 | 0.0245 |
| Late (23-32) | 1.24 ± 0.48 (7) | 2.68 ± 0.79 (10) | 1.51 ± 0.46 (7) | 0.2691 | 0.0945 | 0.7029 | 0.1378 |
|  |  |  |  | P>0.05 | P<0.05 | P<0.01 | P<0.001 |

### EGFR/ERBB3-dependent cell survival is compromised in GM3 synthase deficiency cells

EGFR and ERBB3 signaling support cell survival [37, 38]. Fluctuations in these signaling pathways may therefore lead to decreased cell survival of differentiating cells. We investigated the impact of altered abundance of these essential RTKs on cell survival by assessing the appearance of cleaved caspase3, an apoptotic effector, in differentiating GM3SD cells (Figure 11). While cleaved caspase3 was detected in WT and both GM3SD iPSCs, it decreased to undetectable levels in WT cells by the mid-point of differentiation to NCCs (Figure 11A). In contrast, levels of cleaved caspase3 remained detectable through d13 in differentiating p.Glu355Lys cells (Figure 11B) and through d17 in p.Arg288Ter cells (Figure 11C). This result suggested that GM3SD cells are more prone to programmed cell death during differentiation to NCCs, a phenotype that may relate to altered EGFR/ErbB3 abundance.

**Figure 11.**
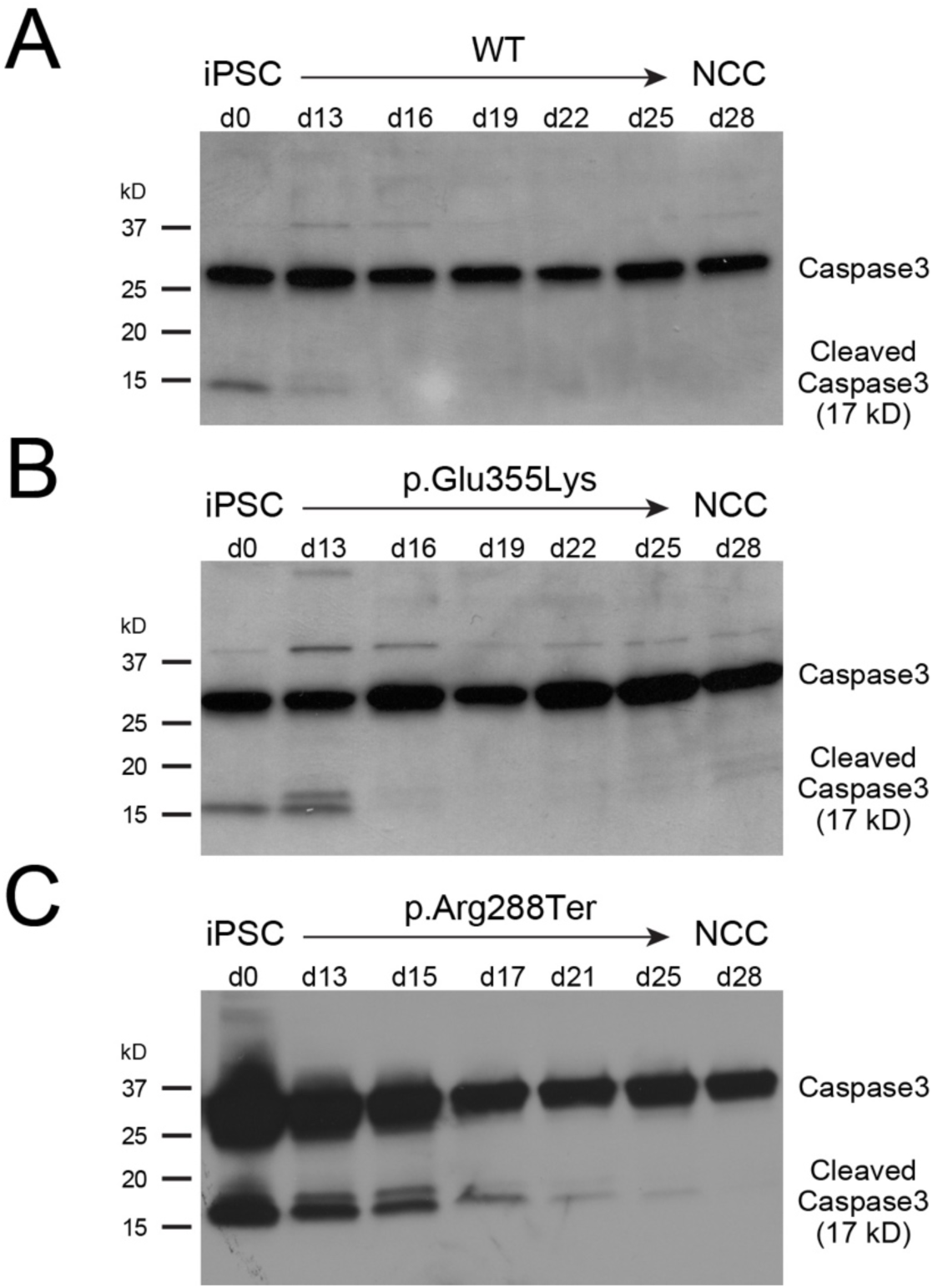
Apoptosis is enhanced in GM3SD cells during differentiation to NCCs. Apoptosis, reported by the abundance of cleaved Caspase3, is detectable in WT and GM3SD iPSC cultures (**A,B,C**) at day 0 (***d0***). As WT cells differentiate to NCCs (**A**), cleaved Caspase3 is barely detectable at the midpoint of the differentiation course (***d13***), whereas it remains at or above iPSC levels in p.Glu355Lys (**B**) and p.Arg288Ter (**C**) cells at the same time point and beyond.

In order to test the hypothesis that ganglioside deficiency increases the likelihood of cell death, we stressed the EGFR/ERBB3 signaling pathway by exposing cells to erlotinib, a pharmacologic inhibitor of EGFR signaling, at the midpoint (d11 – d13) of iPSC differentiation to NCCs [39]. We observed that treatment with erlotnib resulted in significant detachment of presumably apoptotic cells from the surface of the tissue culture well. GM3SD cells dissociated from the surface of the culture well to a much greater extent than WT cells and significant levels of cleaved caspase3 were detected within the floating cell population (Figure 12A). Exposure of WT cells to 100 µM erlotinib for 24 - 48 hours resulted in a consistent and time-dependent increase in the ratio of cleaved to full-length caspase3, although the increase did not achieve statistical signifcance (Table 3 and Figure 12B). In both populations of GM3SD cells, cleaved caspase3 levels were markedly more variable than in WT with or without erlotnib treatment. Upon erlotnib challenge, p.Glu355Lys and p.Arg288Ter cells exhibited statistically significant increases in cleaved caspase3 at one or more timepoints whether the GM3SD cells were compared to erlotnib-treated or vehicle-treated WT cells (Table 3 **and** Figure 12B). Collectively, these findings demonstrate an increased sensitivity of the GM3SD cells to erlotinib during NCC differentiation, a phenotype that may relate to unregulated expression/abundance of RTKs such as EGFR/ERBB3 during critical stages of differentiation.

**Figure 12.**
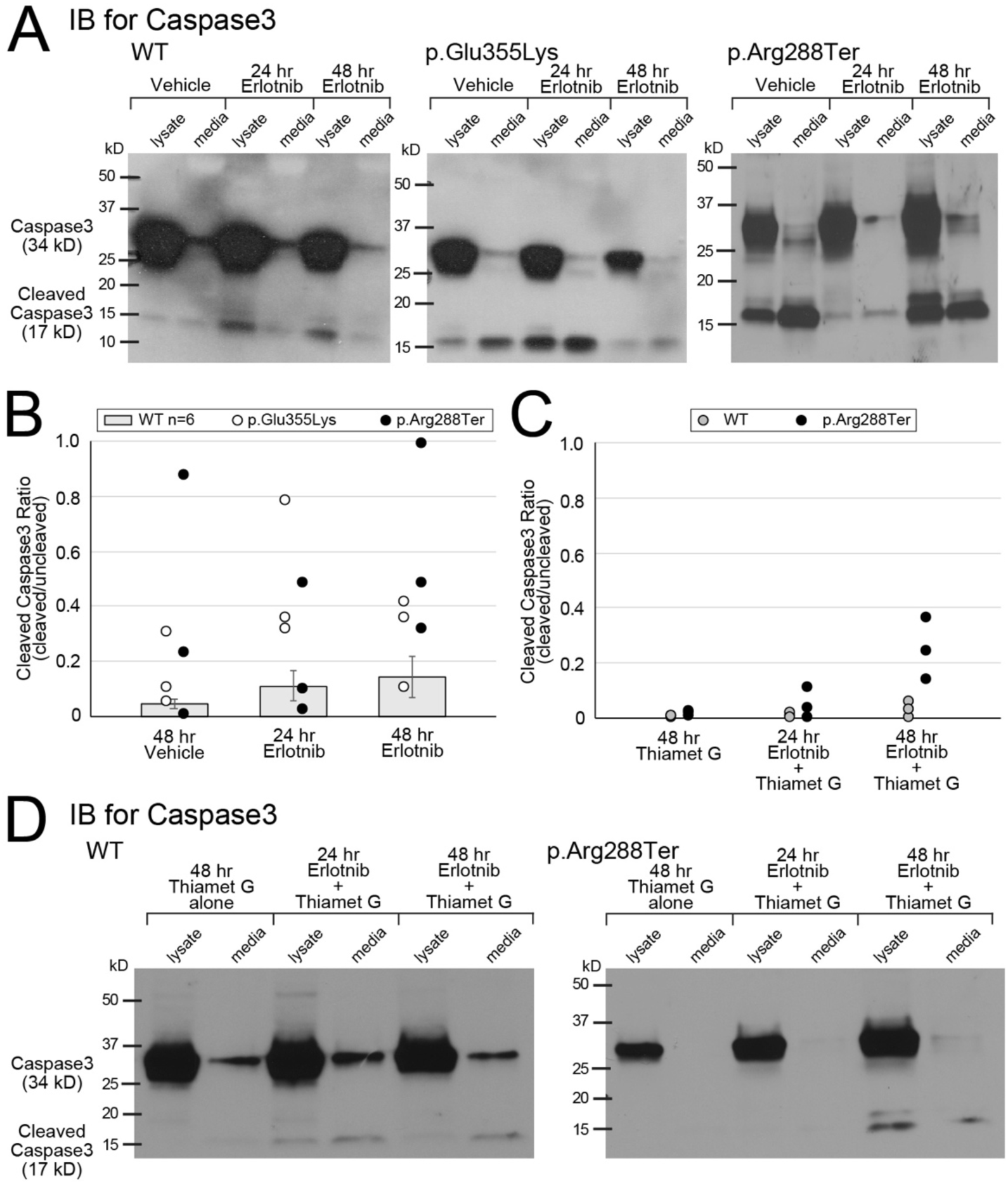
Pharmacologic intervention reveals sensitivity of GM3SD cells to altered EGFR signaling and rescue of apoptosis by increased protein O-GlcNAcylation. (**A**) Treatment of WT or GM3SD cells with erlotnib, an EGFR antagonist, during mid-differentiation to NCC (d11-14) induces increased Caspase3 cleavage. Caspase3 is detected in the cell lysate (***lysate***) of plated cells and in the cells recovered from the media (***media***) by centrifugation (quantified in panel B). (**B**) GM3SD cells are more sensitive to erlotnib treatment than WT cells (GM3SD data points present the media and lysate combined for 3 independent differentiations and are plotted on top of bars that present mean ± SEM for n=6 independent differentiations of WT cells treated similarly). (**C**) Thiamet G treatment, which enhances protein O-GlcNAcylation by inhibition of O-GlcNAcase, decreases Caspase3 cleavage in WT and GM3SD cells. (**D**) Representative Caspase3 western blots demonstrate reduction in Caspase3 cleavage in presence of thiamet G (quantified in panel C).

**Table 3.**
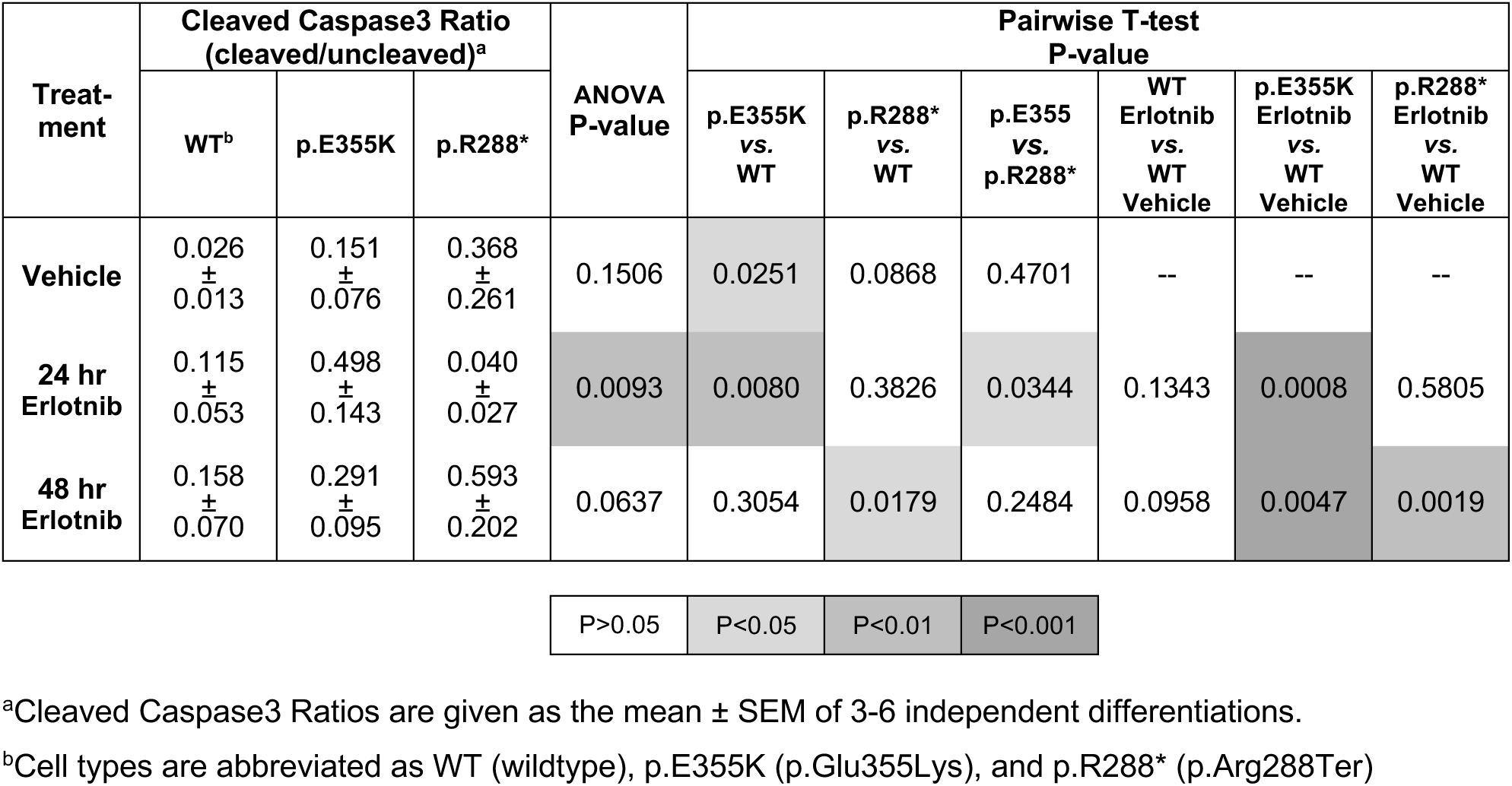
Cleaved Caspase3 in wildtype and GM3SD variant cells at mid-differentiation with and without EGFR inhibition by erlotnib.

| Treat-<br>ment | Cleaved Caspase3 Ratio<br>(cleaved/uncleaved)<sup>a</sup> |  |  | ANOVA<br>P-value | Pairwise T-test<br>P-value |  |  |  |  |  |
| --- | --- | --- | --- | --- | --- | --- | --- | --- | --- | --- |
|  | WT<sup>b</sup> | p.E355K | p.R288* |  | p.E355K<br>vs.<br>WT | p.R288*<br>vs.<br>WT | p.E355<br>vs.<br>p.R288* | WT<br>Erlotinib<br>vs.<br>WT<br>Vehicle | p.E355K<br>Erlotinib<br>vs.<br>WT<br>Vehicle | p.R288*<br>Erlotinib<br>vs.<br>WT<br>Vehicle |
| **Vehicle** | 0.026<br>±<br>0.013 | 0.151<br>±<br>0.076 | 0.368<br>±<br>0.261 | 0.1506 | 0.0251 | 0.0868 | 0.4701 | -- | -- | -- |
| **24 hr**<br>**Erlotinib** | 0.115<br>±<br>0.053 | 0.498<br>±<br>0.143 | 0.040<br>±<br>0.027 | 0.0093 | 0.0080 | 0.3826 | 0.0344 | 0.1343 | 0.0008 | 0.5805 |
| **48 hr**<br>**Erlotinib** | 0.158<br>±<br>0.070 | 0.291<br>±<br>0.095 | 0.593<br>±<br>0.202 | 0.0637 | 0.3054 | 0.0179 | 0.2484 | 0.0958 | 0.0047 | 0.0019 |

|  |  |  |  |
| --- | --- | --- | --- |
| P>0.05 | P<0.05 | P<0.01 | P<0.001 |

<sup>a</sup>Cleaved Caspase3 Ratios are given as the mean ± SEM of 3-6 independent differentiations.
<sup>b</sup>Cell types are abbreviated as WT (wildtype), p.E355K (p.Glu355Lys), and p.R288\* (p.Arg288Ter)

### Pharmacologic inhibition of O-GlcNAcase rescues viability of GM3 synthase deficiency cells

Protein O-GlcNAcylation is responsive to many cellular response pathways and has been proposed to serve as a signal integrator that regulates downstream effectors [40]. The increase in O-GlcNAcylation that we detected in WT NCCs compared to iPSCs and the indication that absolute levels of protein O-GlcNAcylation are more variable in GM3SD cells than in WT (see Figure 8), suggested that pharmacologic manipulation of O-GlcNAcylation might attenuate the impact of altered EGFR/ERBB3 expression in GM3SD cells. To test this hypothesis, we treated WT and p.Arg288Ter cells with thiamet G, a specific inhibitor of the hexosaminidase (O-GlcNAcase) responsible for removing O-GlcNAc from nuclear and cytoplasmic proteins, with or without co-treatment with erlotnib (Figure 12C,D). While thiamet G treatment tended to decrease cleaved caspase3 in WT cells, the decreases were not significant when comparing thiamet G treatment to vehicle alone, erlotnib alone, or to thiamet G + erlotnib treatment at 24 or 48 hours of exposure (Table 4). Dual treatment of p.Arg288Ter cells with erlotnib and thiamet G resulted in cleaved caspase3 levels that were statistically indistinguishable from erlotnib treated WT cells at 24 and 48 hours of exposure, indicating that increased O-GlcNAcylation rescued erlotnib-induced apoptosis of GM3SD cells to WT levels (Figure 12C,D **and** Table 4).

**Table 4.** Cleaved Caspase3 in wildtype and GM3SD variant cells at mid-differentiation in presence of EGFR (erlotnib) and O-GlcNAcase (thiamet G) inhibition.

| Treatment | Cleaved Caspase3 Ratio<br>(cleaved/uncleaved)<sup>a</sup> |  | Pairwise T-test<br>P-value |  |  |  |
| --- | --- | --- | --- | --- | --- | --- |
|  | Wildtype | p.Arg288Ter | Wildtype<br>Thiamet G<br>vs.<br>Wildtype<br>Vehicle<sup>b</sup> | Wildtype<br>Erlotinib +<br>Thiamet G<br>vs.<br>Wildtype<br>Thiamet G | Wildtype<br>Erlotinib<sup>b</sup><br>vs.<br>Wildtype<br>Erlotinib +<br>Thiamet G | p.Arg288Ter<br>Erlotinib +<br>Thiamet G<br>vs.<br>Wildtype<br>Erlotinib<sup>b</sup> |
| **Vehicle<sup>b</sup>** | 0.026 ± 0.013 | 0.368 ± 0.261 | 0.2455 | -- | -- | -- |
| **Thiamet G**<br>**alone** | 0.002 ± 0.001 | 0.012 ± 0.005 | -- | -- | -- | -- |
| **24 hr**<br>**Erlotinib +**<br>**Thiamet G** | 0.016 ± 0.007 | 0.048 ± 0.031 | -- | 0.1101 | 0.2462 | 0.4326 |
| **48 hr**<br>**Erlotinib +**<br>**Thiamet G** | 0.028 ± 0.016 | 0.245 ± 0.064 | -- | 0.1911 | 0.2531 | 0.4569 |

P>0.05
<sup>a</sup>Cleaved Caspase3 Ratios are given as the mean ± SEM of 3-6 independent differentiations.
<sup>b</sup>Values for these Cleaved Caspase3 Ratios taken from Table 3.

## DISCUSSION

Complex gangliosides are essential components of the extracellular leaflet of the plasma membrane of neural cells. Subsets of these GSLs, including GM3, preferentially partition into lipid microdomains that are also enriched in cell-signaling receptors and adhesion molecules [30, 31]. Therefore, mutations in the biosynthetic enzymes responsible for generating GSLs would be expected to impart significant pathology. Surprisingly, however, the first reports of a mouse knockout of *St3gal5* described a relatively normal animal capable of surviving and reproducing [6]. Subsequent analysis demonstrated more subtle defects such as cochlear neuronal cell death, reduced male fertility, altered insulin receptor signaling, and skewed inflammatory responses [6, 41–44]. The relatively mild nature of the mouse ST3Gal5 knockout phenotype has been proposed to result from the ability of that organism to efficiently extend and sialylate LacCer on external Gal residues to generate o-series GSLs (Figure 1). These gangliosides, which we refer to here as alternatively sialylated gangliosides, were proposed to be capable of at least partially compensating for loss of the a-, b-, and c-series gangliosides [45]. Our previously published work on p.Glu355Lys patient fibroblasts demonstrated that human fibroblasts do not possess a robust capacity to produce significant amounts of o-series, alternatively sialylated gangliosides [8]. It was thought that in the absence of this alternative GSL biosynthetic capacity, patients with GM3SD manifest significant neurologic complications that are not currently well modeled in the mouse. Therefore, we took advantage of iPSC technology to investigate the cellular consequences of altered ganglioside biosynthesis during human neural cell differentiation in two different GM3SD variants.

Consistent with our previous report on p.Glu355Lys fibroblasts, GSL profiles of iPSCs and NCCs from both GM3SD variants were devoid of GM3 and GM3-derived GSLs [8, 10]. Other than loss of GM3, the GSL profile of GM3SD iPSCs was strikingly similar to WT, suggesting that GM3SD should be expected to have a minimal impact on the maintenance of pluripotency and, perhaps, of neural stem cell populations in developing or mature brains. However, the value of modeling human disease in iPSC-derived cell populations became apparent upon differentiation of GM3SD cells to NCC, as neural-specific and variant-specific changes in GSL profiles became evident. In GM3SD NCCs, the absence of a-, b-, and c-series gangliosides was associated with large increases in LacCer, the precursor to GM3, and parallel increases in globo- and lacto-series GSLs compared to WT. Other neural-specific changes in GM3SD cells included enhanced production of the alternatively sialylated gangliosides GM1b and GD1c in p.Glu355Lys and greater retention of LacCer in p.Arg288Ter cells compared to each other and to WT. Another more subtle difference between the GSL profiles of GM3SD cells was noted in the detection and characterization of minor GSLs with extended LacNAc structures. Non-sialylated extended LacNAc structures were more abundant in p.Arg288Ter NCCs, whereas extended LacNAc structures were more likely to be sialylated in p.Glu355Lys NCCs. Thus, a hallmark of human GM3SD is the retention and partial rerouting of LacCer into other biosynthetic pathways. The extent to which the clinical phenotypes of GM3SD arise from elevated LacCer, reduced complex ganglioside biosynthesis, or a combination of both remains unknown, but the iPSCs and NCCs we describe here provide a useful platform to investigate the impact of GSL imbalance at the cellular level.

It is important to note that neither the p.Glu355Lys or p.Arg288Ter variant polypeptides have yet been detected as endogenously expressed proteins in human tissues or derived cells. However, both proteins are produced when expressed as epitope-tagged forms in HEK cells but lack measurable transferase activity [9]. Thus, while both variants are functionally null, it remains to be determined whether mRNA produced from an endogenous variant *ST3GAL5* gene locus can lead to the translation of a stable enzyme variant. The molecular nature of the two ST3GAL5 protein variants fails to suggest an obvious mechanism by which either might differentially impact alternative sialylation. Thus, the mechanism for the retention of LacCer precursor or for the production of alternatively sialylated GSLs most likely entails a nuanced, but yet-to-be characterized, shift in the functional organization of GSL processing in the secretory pathway. Such a reorganization might arise from altered cell-surface signaling or from the association of non-functional ST3Gal5 protein with other glycosyltransferases. The latter of these non-exclusive possibilities has been proposed as a general feature of enzymes involved in GSL biosynthesis [46, 47].

SEEL-based capture of cell surface glycoproteins in WT and p.Glu355Lys NCCs identified both enrichment and loss of membrane raft associated proteins whose GO functions indicate roles in cell adhesion, cell signaling, axon pathfinding, neurogenesis, nervous system development, and cell migration. Neuropathological data at cellular resolution is not yet available for GM3SD brain tissue, making it difficult to associate specific cellular and developmental processes (neuroblast/neural stem cell proliferation and migration, axonal/dendritic outgrowth, neuropil expansion, myelination, etc.) with loss of ganglioside expression. However, delayed neurologic development and reduced cranial growth are characteristic of GM3SD and point to likely impairments in the types of functions associated with the proteins whose presence we detected as altered at the cell surface of GM3SD cells [1]. Minimally, these changes are consistent with material changes in the physicochemical nature of the plasma membrane and in membrane subdomains relevant for signaling and adhesion.

Although the molecular mechanisms by which specific GSLs contribute to the self-associative generation or stabilization of signaling microdomains remain elusive, direct interactions between GM3 and RTKs such as EGFR have been functionally and structurally demonstrated [33, 48–51]. Weak, potentially disruptive binding between EGFR or other RTKs and the alternatively sialylated GSLs produced by GM3SD cells may significantly impact the efficiency of receptor partition into signaling domains, the efficacy of their signaling activity, or the extent of their residence time at the cell surface (Figure 13). As pluripotent WT cells proceed down any particular differentiation pathway (e.g. to NCCs), receptor signaling activates and inhibits various cellular effector functions to appropriately tune or balance responses toward achieving a particular endpoint. Phosphorylation and O-GlcNAcylation of effector proteins are frequently responsible for driving these responses (Figure 13A). We propose that GM3SD generates aberrant populations of signaling domains that are more heterogeneous in their size and/or functionality compared to WT (Figure 13B). Some domains may be too small or unstable to accommodate the signaling machinery of specific receptors while others may be larger than in WT, resulting in suppressed or enhanced signaling, respectively, and/or altered residence time for receptors at the surface. Together, the impact of the modified membrane domains results in altered effector responses that impact survivability and other yet-to-be identified characteristics of GM3SD cells.

**Figure 13.**
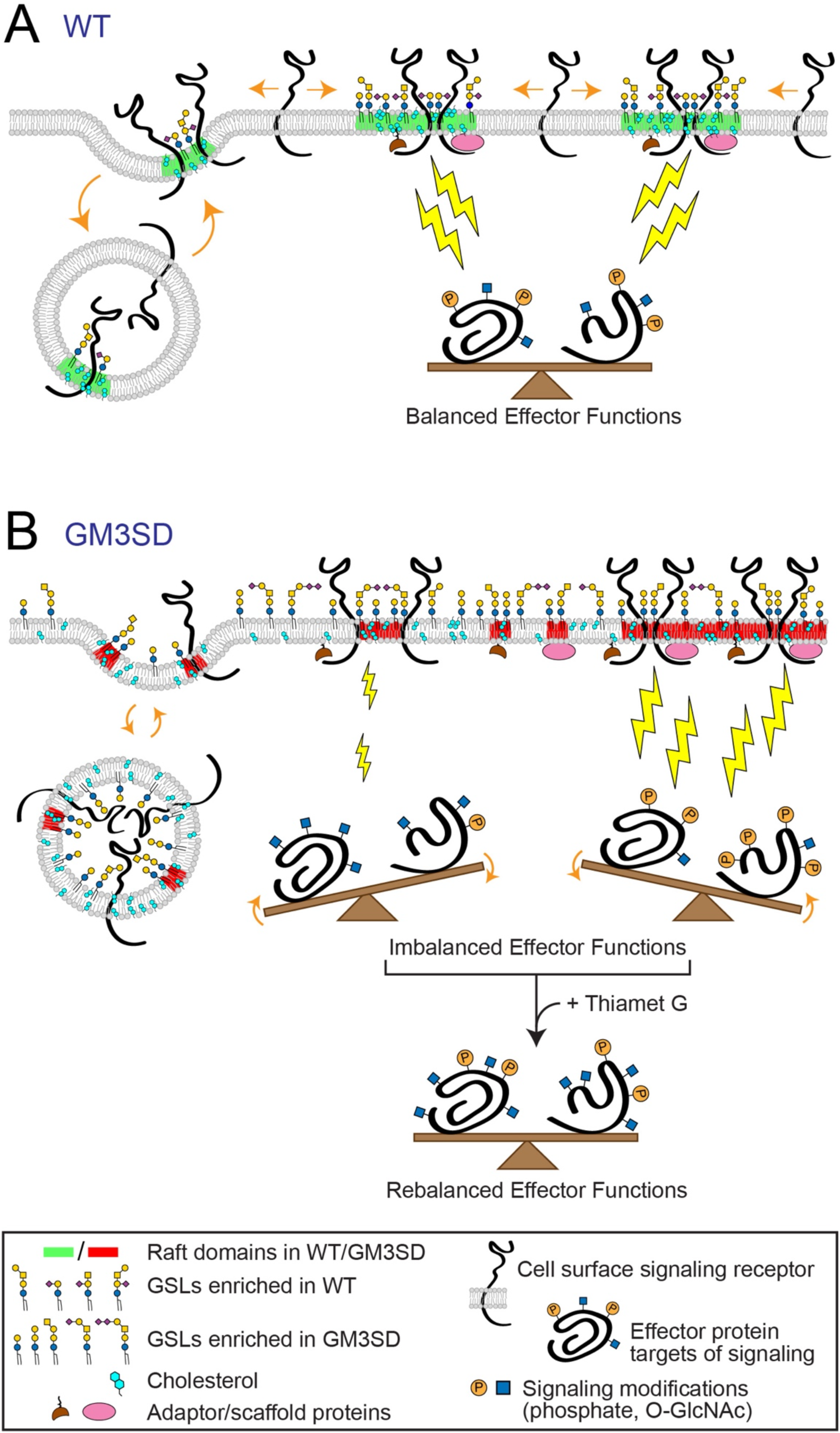
GSL composition impacts lipid rafts and cell signaling in GM3SD. Cell surface signaling receptors are regulated by their migration in-and-out of membrane signaling domains and by the balance of their internalization/reinsertion kinetics. In WT and GM3SD cells, lipid raft characteristics reflect the physicochemical properties of the GSLs, phospholipids, and sterols that constitute the plasma membrane. (**A**) When cells are able to synthesize GM3 and its subsequent end-products, lipid rafts (***shaded green***) facilitate the association of signaling receptors and adaptor/scaffold proteins to appropriately stimulate downstream effectors (***lightning bolts***). Membrane recycling also buffers signaling responses by internalizing surface receptors and shuttling additional signaling capacity from intracellular storage. Effector functions are balanced by appropriate receptor activation, membrane insertion, and physical interactions within signaling domains. (**B**) In the absence of GM3, GSL compositions are shifted toward greater abundance of neutral lipids and toward the appearance of alternatively sialylated species. The characteristics of the resulting lipid rafts (***shaded red***) reflect the ability of these GSLs to interact with receptors and with other raft components. Consequently, increased or decreased signaling through modified rafts may generate imbalanced effector functions that impact cellular responses. Pharmacologic interventions, such as increasing protein O-GlcNAcylation (thiamet G), may establish an alternative balance in effector functions that counters otherwise deleterious downstream processes.

While protein phosphorylation is well recognized as a regulatory post-translational modification, many nuclear and cytoplasmic proteins are also modified by the addition of O-linked GlcNAc at Ser/Thr residues in response to a broad range of cellular signaling activities. O-GlcNAcylation is dynamically driven by the cellular micro-environment, nutrient availability, cell stress, cell cycle, growth factors and associated developmental events, as well as other signals [40]. Thus, like phosphorylation, the profile of O-GlcNAcylated proteins reflects the integration of signals through multiple pathways. We demonstrated that protein O-GlcNAcylation increased as WT and GM3SD iPSCs differentiated to NCCs but the range of O-GlcNAcylation was much broader in GM3SD cells. By pharmacologically enhancing O-GlcNAcylation with thiamet G, cleaved caspase3 levels were significantly reduced in WT and GM3SD cells in the presence or absence of erlotnib [52, 53]. This intervention suggests that, despite broad variation in the expression of primary signaling receptors like EGFR and ERBB3, modulation of downstream effectors can rescue the impact of altered GSL biosynthesis on basic cellular processes in neural lineages (Figure 13B).

An important caveat for the studies reported here is that our observations are based on iPSCs derived from a single representative of each of the p.Glu355Lys and p.Arg288Ter variants. It will be essential to evaluate individual-to-individual variation as more patient samples become available. Nonetheless, individual-to-individual variation in plasma GSL abundance and ceramide diversity was not previously identified as a confounding factor in the p.Arg288Ter GM3SD patient population [54]. Comparing 8 unrelated p.Arg288Ter individuals to the mean value for 19 non-affected individuals in this prior study, the fold-increase in LacCer abundance was 1.9 ± 0.1 (CV%=5.2%) and the fold-increase in the higher mass ceramides of LacCer was 1.6 ± 0.2 (CV%=12.5%). Thus, at least within the p.Arg288Ter population, the major changes in GSL diversity and abundance are well conserved across unrelated individuals. While additional p.Glu355Lys patients are unlikely to become available for future study, expanded efforts to acquire a broader range of p.Arg288Ter and other variant cell types will greatly enhance our understanding of the biochemical corelates of clinical presentation.

Our results demonstrate the importance of investigating glycosylation changes associated with mutations in glycan biosynthetic enzymes in appropriate cell types. We previously described collateral changes in N-linked and O-linked glycosylation of glycoproteins expressed by p.Glu355Lys fibroblasts [8]. Such collateral changes are of interest because ST3GAL5 does not act on glycans linked to proteins; it is absolutely GSL-specific. Thus, the matrix of regulatory processes that control cellular glycosylation are responsive to changes that span glycan classes. This phenomenon has also recently been demonstrated in glyco-engineered human cell lines [55]. The results presented here provide the first characterization of GSL biosynthetic, cell signaling, and cellular survival changes associated with human-derived neural cells in GM3SD. We have identified global changes in GSL biosynthesis that correspond to substantial alterations in cell signaling and residence of glycoproteins at the plasma membrane. We have also identified more subtle changes that may provide differential characteristics capable of distinguishing GM3SD variants. As additional cohorts of GM3SD patients are identified and become available for analysis at the cellular level, the results reported here will provide a framework for assessing functional differences that may provide insight into clinical phenotypes. The iPSC platform we have characterized also presents opportunities to assess the potency of molecular and small-molecule therapeutic interventions that might resolve these cellular phenotypes and impact broader glycomic and pathophysiologic consequences associated with GM3SD.

## Acknowledgements

The authors gratefully acknowledge Huiya Yang and Guangping Gao (University of Massachusetts Medical School) for providing GM3SD p.Arg288Ter fibroblasts and Charles Schwartz (Greenwood Genetics Center) for providing GM3SD p.Glu355Lys fibroblasts. This work was supported by grants from the W.M. Keck Foundation (to M.T., R.S., K.S., and S.D.); National Institutes of Health/National Institute of General Medical Sciences (NIH/NIGMS P41GM103490 to M.T., R.S., and S.D); National Institutes of Health/National Institute of Allergy and Infectious Disease (R21AI129873 to K.A.). M.D. was supported by an NIH/NIGMS T32 Training grant (T32GM107004). The content is solely the responsibility of the authors and does not necessarily represent the official views of the National Institutes of Health.

## REFERENCES

1. Bowser, L.E., et al., Recessive GM3 synthase deficiency: Natural history, biochemistry, and therapeutic frontier. Mol Genet Metab, 2019. 126(4): p. 475–488.

2. Saul, R., G. Wilkes, and R. Stevenson. Salt-And-Pepper’pigmentary changes with severe mental retardation: a new neurocutaneous syndrome. in Proc. Greenwood Genet., Ctr. 1983.

3. Schnaar, R.L., R. Gerardy-Schahn, and H. Hildebrandt, Sialic acids in the brain: gangliosides and polysialic acid in nervous system development, stability, disease, and regeneration. Physiol Rev, 2014. 94(2): p. 461–518.

4. Sipione, S., et al., Gangliosides in the Brain: Physiology, Pathophysiology and Therapeutic Applications. Frontiers in Neuroscience, 2020. 14(1004).

5. Schnaar, R.L., Gangliosides of the Vertebrate Nervous System. J Mol Biol, 2016. 428(16): p. 3325–3336.

6. Yamashita, T., et al., Enhanced insulin sensitivity in mice lacking ganglioside GM3. Proceedings of the National Academy of Sciences, 2003. 100(6): p. 3445–3449.

7. Lloyd, K.O. and K. Furukawa, Biosynthesis and functions of gangliosides: recent advances. Glycoconjugate Journal, 1998. 15(7): p. 627–636.

8. Boccuto, L., et al., A mutation in a ganglioside biosynthetic enzyme, ST3GAL5, results in salt & pepper syndrome, a neurocutaneous disorder with altered glycolipid and glycoprotein glycosylation. Human Molecular Genetics, 2013. 23(2): p. 418–433.

9. Indellicato, R., et al., Total loss of GM3 synthase activity by a normally processed enzyme in a novel variant and in all ST3GAL5 variants reported to cause a distinct congenital disorder of glycosylation. Glycobiology, 2019. 29(3): p. 229–241.

10. Simpson, M.A., et al., Infantile-onset symptomatic epilepsy syndrome caused by a homozygous loss-of-function mutation of GM3 synthase. Nature Genetics, 2004. 36(11): p. 1225–1229.

11. Fragaki, K., et al., Refractory epilepsy and mitochondrial dysfunction due to GM3 synthase deficiency. European Journal of Human Genetics, 2013. 21(5): p. 528–534.

12. Gordon-Lipkin, E., et al., ST3GAL5-Related Disorders: A Deficiency in Ganglioside Metabolism and a Genetic Cause of Intellectual Disability and Choreoathetosis. J Child Neurol, 2018. 33(13): p. 825–831.

13. Lee, J.S., et al., GM3 synthase deficiency due to ST3GAL5 variants in two Korean female siblings: Masquerading as Rett syndrome-like phenotype. American Journal of Medical Genetics Part A, 2016. 170(8): p. 2200–2205.

14. Li, T.A. and R.L. Schnaar, Chapter Two - Congenital Disorders of Ganglioside Biosynthesis, in Progress in Molecular Biology and Translational Science, R.L. Schnaar and P.H.H. Lopez, Editors. 2018, Academic Press. p. 63–82.

15. Si-Tayeb, K., et al., Generation of human induced pluripotent stem cells by simple transient transfection of plasmid DNA encoding reprogramming factors. BMC Dev Biol, 2010. 10: p. 81.

16. Menendez, L., et al., Directed differentiation of human pluripotent cells to neural crest stem cells. Nat Protoc, 2013. 8(1): p. 203–12.

17. Cederquist, G.Y., et al., A Multiplex Human Pluripotent Stem Cell Platform Defines Molecular and Functional Subclasses of Autism-Related Genes. Cell Stem Cell, 2020. 27(1): p. 35–49.e6.

18. Aoki, K., et al., Dynamic developmental elaboration of N-linked glycan complexity in the Drosophila melanogaster embryo. J Biol Chem, 2007. 282(12): p. 9127–42.

19. Nimrichter, L., et al., E-selectin receptors on human leukocytes. Blood, 2008. 112(9): p. 3744–52.

20. Anumula, K.R. and P.B. Taylor, A comprehensive procedure for preparation of partially methylated alditol acetates from glycoprotein carbohydrates. Anal Biochem, 1992. 203(1): p. 101–8.

21. Vukelić, Ž., et al., Screening and sequencing of complex sialylated and sulfated glycosphingolipid mixtures by negative ion electrospray Fourier transform ion cyclotron resonance mass spectrometry. Journal of the American Society for Mass Spectrometry, 2005. 16(4): p. 571–580.

22. Varki, A., et al., Symbol Nomenclature for Graphical Representations of Glycans. Glycobiology, 2015. 25(12): p. 1323–4.

23. Wells, L., G.W. Hart, and D. Athens Guidelines for the Publication of Glycomics, Glycomics: building upon proteomics to advance glycosciences. Mol Cell Proteomics, 2013. 12(4): p. 833–5.

24. York, W.S., et al., MIRAGE: the minimum information required for a glycomics experiment. Glycobiology, 2014. 24(5): p. 402–6.

25. Rojas-Macias, M.A., et al., Towards a standardized bioinformatics infrastructure for N- and O-glycomics. Nat Commun, 2019. 10(1): p. 3275.

26. Mbua, N.E., et al., Selective Exo-Enzymatic Labeling of N-Glycans on the Surface of Living Cells by Recombinant ST6Gal I. Angewandte Chemie International Edition, 2013. 52(49): p. 13012–13015.

27. Sun, T., et al., One-Step Selective Exoenzymatic Labeling (SEEL) Strategy for the Biotinylation and Identification of Glycoproteins of Living Cells. Journal of the American Chemical Society, 2016. 138(36): p. 11575–11582.

28. Meng, L., et al., Enzymatic basis for N-glycan sialylation: structure of rat α2,6-sialyltransferase (ST6GAL1) reveals conserved and unique features for glycan sialylation. J Biol Chem, 2013. 288(48): p. 34680–98.

29. Brown, D.A. and E. London, Structure of detergent-resistant membrane domains: does phase separation occur in biological membranes? Biochem Biophys Res Commun, 1997. 240(1): p. 1–7.

30. Brown, D.A. and J.K. Rose, Sorting of GPI-anchored proteins to glycolipid-enriched membrane subdomains during transport to the apical cell surface. Cell, 1992. 68(3): p. 533–44.

31. Sandhoff, K. and T. Kolter, Biosynthesis and degradation of mammalian glycosphingolipids. Philosophical transactions of the Royal Society of London. Series B, Biological sciences, 2003. 358(1433): p. 847–861.

32. Simons, K. and E. Ikonen, Functional rafts in cell membranes. Nature, 1997. 387(6633): p. 569–72.

33. Miljan, E.A., et al., Interaction of the Extracellular Domain of the Epidermal Growth Factor Receptor with Gangliosides. Journal of Biological Chemistry, 2002. 277(12): p. 10108–10113.

34. Shih, A., S. Telesco, and R. Radhakrishnan, Analysis of Somatic Mutations in Cancer: Molecular Mechanisms of Activation in the ErbB Family of Receptor Tyrosine Kinases. Cancers, 2011. 3: p. 1195–1231.

35. Black, L.E., J.F. Longo, and S.L. Carroll, Mechanisms of Receptor Tyrosine-Protein Kinase ErbB-3 (ERBB3) Action in Human Neoplasia. The American Journal of Pathology, 2019. 189(10): p. 1898–1912.

36. Steinkamp, M.P., et al., erbB3 Is an Active Tyrosine Kinase Capable of Homo- and Heterointeractions. Molecular and Cellular Biology, 2014. 34(6): p. 965–977.

37. Wee, P. and Z. Wang, Epidermal Growth Factor Receptor Cell Proliferation Signaling Pathways. Cancers (Basel), 2017. 9(5).

38. Holbro, T., et al., The ErbB2/ErbB3 heterodimer functions as an oncogenic unit: ErbB2 requires ErbB3 to drive breast tumor cell proliferation. Proceedings of the National Academy of Sciences, 2003. 100(15): p. 8933–8938.

39. Rukazenkov, Y., et al., Epidermal growth factor receptor tyrosine kinase inhibitors: similar but different? Anticancer Drugs, 2009. 20(10): p. 856–66.

40. Hart, G.W., Nutrient regulation of signaling and transcription. J Biol Chem, 2019. 294(7): p. 2211–2231.

41. Yoshikawa, M., et al., Ganglioside GM3 is essential for the structural integrity and function of cochlear hair cells. Human Molecular Genetics, 2015. 24(10): p. 2796–2807.

42. Yoshikawa, M., et al., Mice lacking ganglioside GM3 synthase exhibit complete hearing loss due to selective degeneration of the organ of Corti. Proceedings of the National Academy of Sciences, 2009. 106(23): p. 9483–9488.

43. Sandhoff, R., et al., Novel Class of Glycosphingolipids Involved in Male Fertility *. Journal of Biological Chemistry, 2005. 280(29): p. 27310–27318.

44. Yamashita, T., et al., Interruption of ganglioside synthesis produces central nervous system degeneration and altered axon–glial interactions. Proceedings of the National Academy of Sciences of the United States of America, 2005. 102(8): p. 2725–2730.

45. Collins, B.E., et al., Enhanced Binding of the Neural Siglecs, Myelin-associated Glycoprotein and Schwann Cell Myelin Protein, to Chol-1 (α-Series) Gangliosides and Novel Sulfated Chol-1 Analogs*. Journal of Biological Chemistry, 1999. 274(53): p. 37637–37643.

46. Maccioni, H.J.F., R. Quiroga, and M.L. Ferrari, Cellular and molecular biology of glycosphingolipid glycosylation. Journal of Neurochemistry, 2011. 117(4): p. 589–602.

47. Maccioni, H.J.F., R. Quiroga, and W. Spessott, Organization of the synthesis of glycolipid oligosaccharides in the Golgi complex. FEBS Letters, 2011. 585(11): p. 1691–1698.

48. Bremer, E.G., et al., Ganglioside-mediated modulation of cell growth, growth factor binding, and receptor phosphorylation. J Biol Chem, 1984. 259(11): p. 6818–25.

49. Bremer, E.G., J. Schlessinger, and S. Hakomori, Ganglioside-mediated modulation of cell growth. Specific effects of GM3 on tyrosine phosphorylation of the epidermal growth factor receptor. J Biol Chem, 1986. 261(5): p. 2434–40.

50. Laine, R.A. and S. Hakomori, Incorporation of exogenous glycosphingolipids in plasma membranes of cultured hamster cells and concurrent change of growth behavior. Biochem Biophys Res Commun, 1973. 54(3): p. 1039–45.

51. Wang, X.-Q., P. Sun, and A.S. Paller, Ganglioside GM3 Blocks the Activation of Epidermal Growth Factor Receptor Induced by Integrin at Specific Tyrosine Sites *. Journal of Biological Chemistry, 2003. 278(49): p. 48770–48778.

52. Jiang, M., et al., XBP1 (X-Box-Binding Protein-1)-Dependent O-GlcNAcylation Is Neuroprotective in Ischemic Stroke in Young Mice and Its Impairment in Aged Mice Is Rescued by Thiamet-G. Stroke, 2017. 48(6): p. 1646–1654.

53. Yuzwa, S.A., et al., A potent mechanism-inspired O-GlcNAcase inhibitor that blocks phosphorylation of tau in vivo. Nat Chem Biol, 2008. 4(8): p. 483–90.

54. Aoki, K., et al., Mass spectrometric quantification of plasma glycosphingolipids in human GM3 ganglioside deficiency. Clin Mass Spec, 2019. 14: p. 106–114.

55. Huang, Y.F., et al., Global mapping of glycosylation pathways in human-derived cells. Dev Cell, 2021. 56(8): p. 1195–1209 e7.

