## Supplement Figures for "Neural-specific alterations in glycosphingolipid biosynthesis and cell signaling associated with two human ganglioside GM3 Synthase Deficiency variants"

**SUPPLEMENTAL DATA**

**Supplement Figure 1. Generation and initial characterization of p.Arg288Ter iPS Cells.** Fibroblasts harvested from a p.Arg288Ter patient were reprogrammed to iPS cells as described in Methods. The morphology and expression of pluripotency markers in three independent reprogrammed clones (2.1a, 2.2c, and 2.3j) are shown. All three clones possess the potential to differentiate to neuroectoderm as detected by PAX6 expression.

**Supplement Figure 2. Immunofluorescence staining for pluripotency and neural crest markers in WT and p.Arg288Ter cells.** iPSCs derived from p.Arg288Ter patient fibroblasts express appropriate pluripotency markers (Oct3/4, SOX2, SSEA4). Upon differentiation to neural crest cells, expression of pluripotency markers is lost as expression of neural crest markers (AP2, HNK1, p75) is acquired.

**Supplement Figure 3. Immunofluorescence staining for pluripotency and neural crest markers in WT and p.Glu355Lys cells.** iPSCs derived from p.Glu355Lys patient fibroblasts express appropriate pluripotency markers (Oct3/4, SOX2, SSEA4). Like the p.Arg288Ter iPSCs (Supplement Figure 1), p.Glu355Lys iPSCs are capable of differentiation to neuroectoderm (PAX6 positive) and can also acquire neural crest markers (HNK1, p75) concurrent with loss of pluripotentcy markers.

**Supplement Figure 4. MS2 fragmentation supporting structural assignment of unique glycosphingolipids.** NSI-MS/MS analysis of LacNAc-extended GSLs expressed by GM3SD NCCs. Top panel shows fragmentation of sialylated GSL detected at greater abundance in p.Glu355Lys NCCS than in p.Arg288Ter NCCs. Bottom panel shows fragmentation of non-sialylated GSL detected at greater abundance in p.Arg288Ter NCCs than in p.Arg288TerNCCs.

**Supplement Table 1. Excel spreadsheet summarizing SEEL proteomic data.** The spreadsheet file summarizes the proteins identified through SEEL capture. ***Tab 1***, labelled "All Proteins," presents all of the proteins identified in the SEEL proteomic analysis of differentiating wildtype and p.Glu355Lys cells, regardless of their subcellular localization or number of spectral counts (# PSM, protein spectral match). ***Tab 2***, labelled "Excluded Proteins," presents the small number of non-cell surface proteins identified across all of the SEEL capture experiments. ***Tab 3***, labelled "Non-excluded Proteins > 10 PSM," presents the proteins identified by more than 10 spectral counts and that were not excluded because they are not cell surface proteins (listed in ***Tab 2***). Column X in this Tab presents the ratio change in cell surface proteins in p.Glu355Lys cells compared to wildtype. ***Tab 4***, labelled "GO--Raft and Endosome Proteins," presents proteins that have genome ontology notations consistent with proteins found in lipid rafts or in endosomal recycling pathways.

Supplement Figure 1

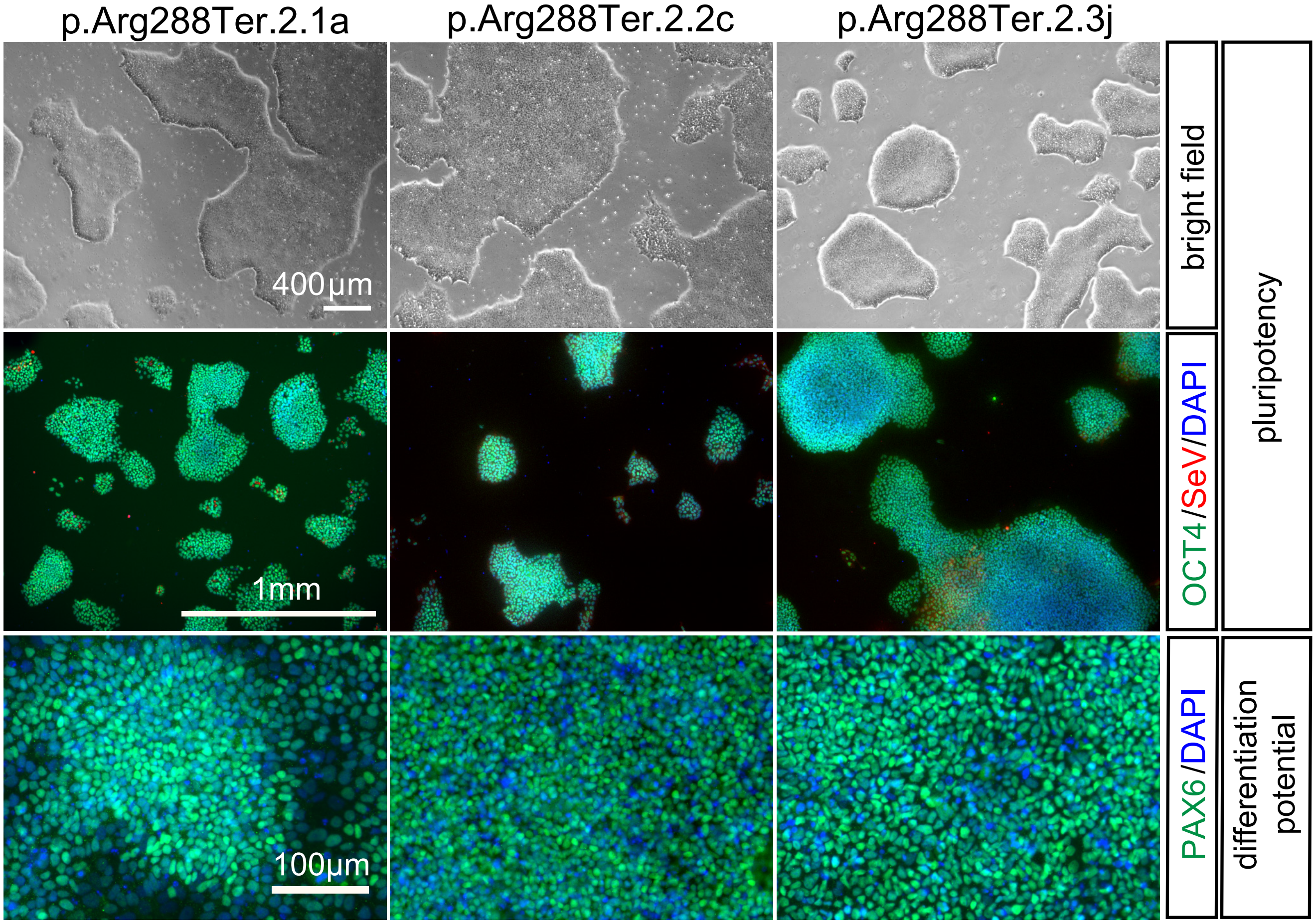

Supplement Figure 2

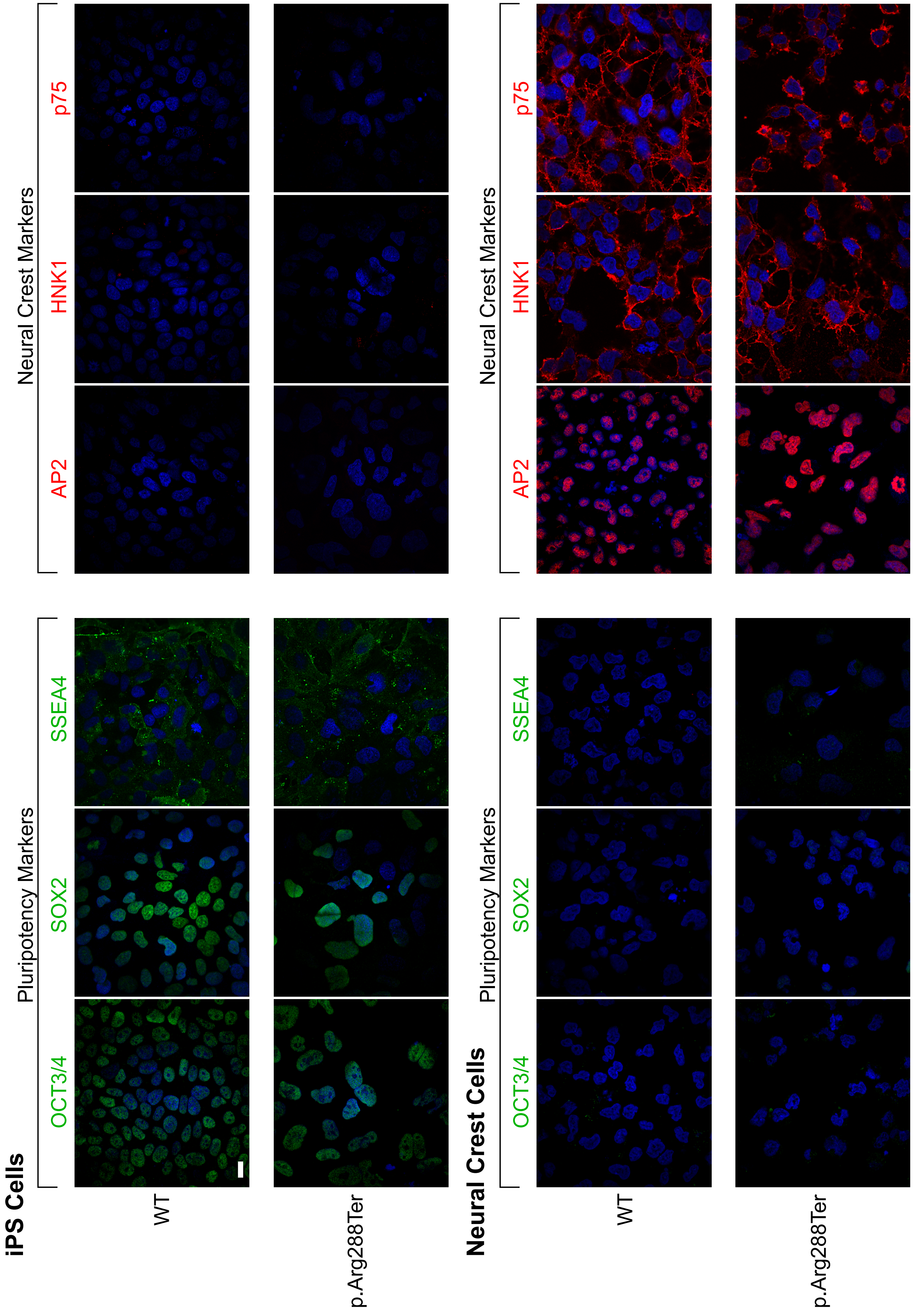

Supplement Figure 3

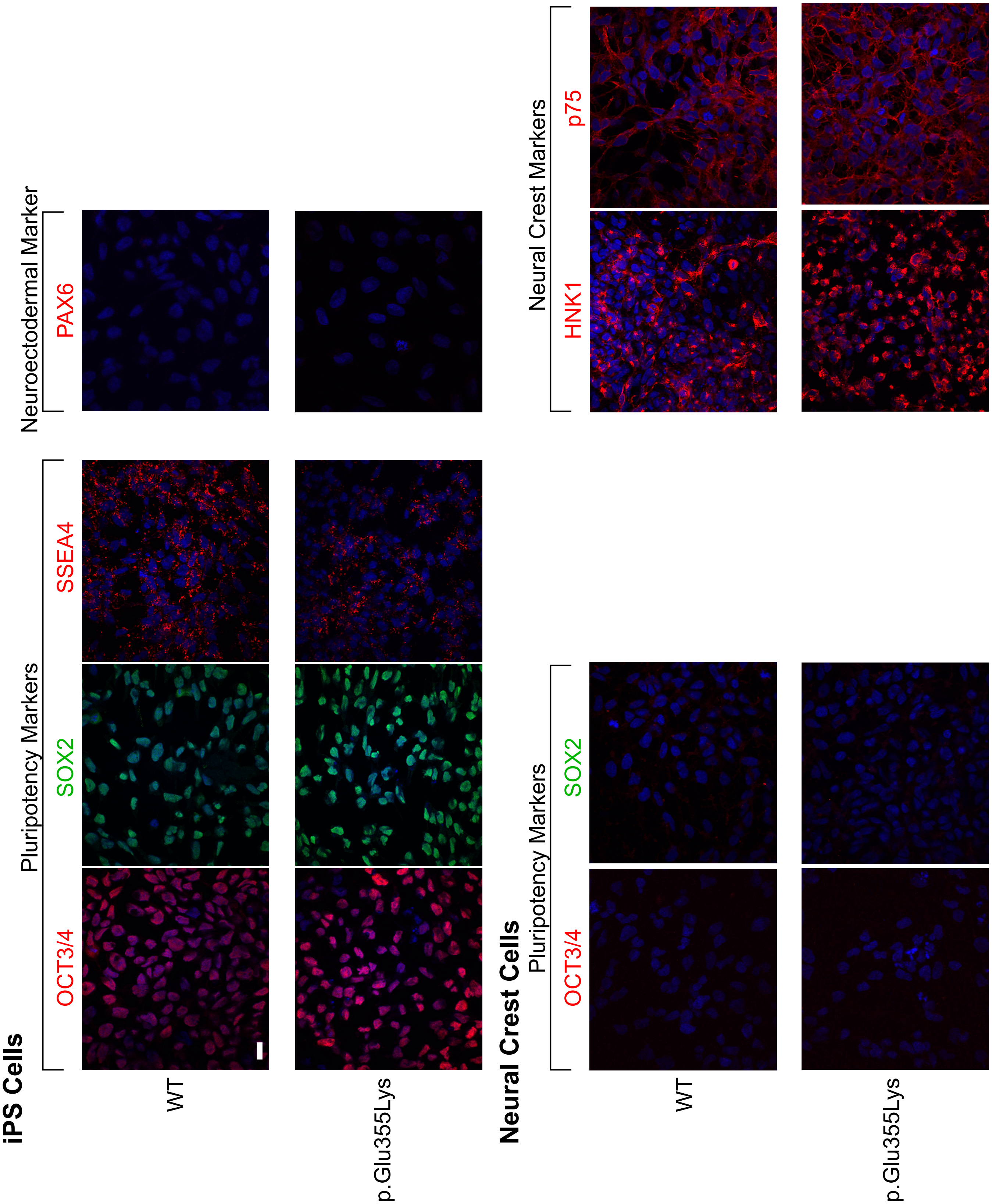

Supplement Figure 4

A

T: ITMS + c NSI Full ms2 1146

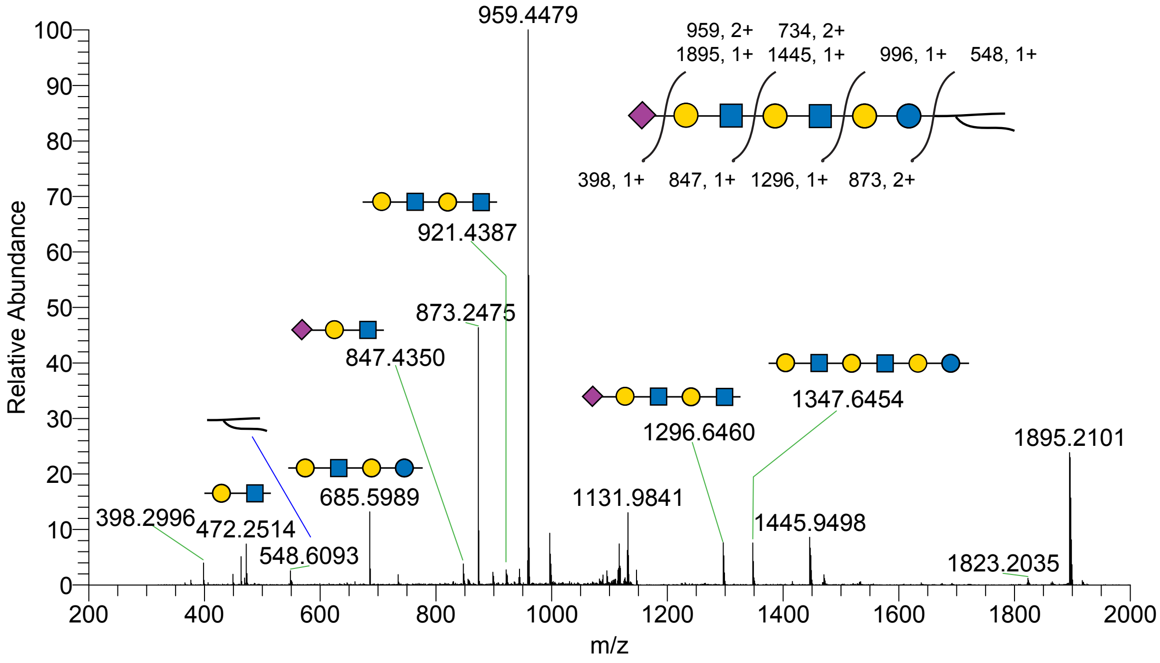


B

T: ITMS + c NSI Full ms2 1192

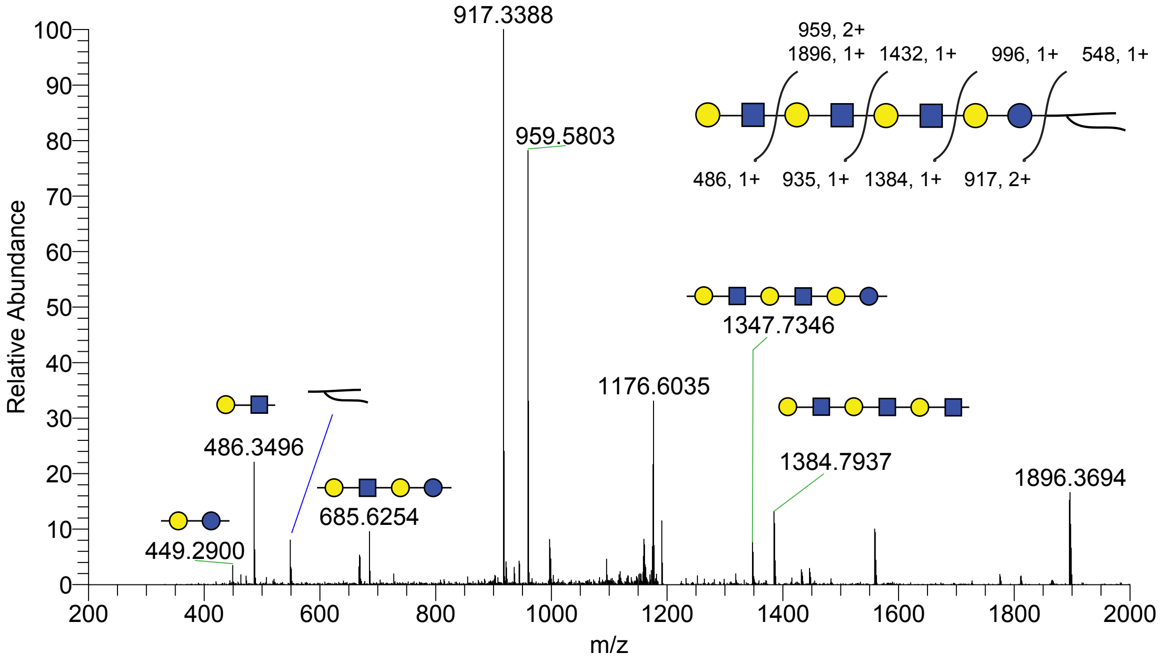
